# Single-cell multi-omics decodes regulatory programs during development of mouse secondary palate

**DOI:** 10.1101/2022.11.02.514609

**Authors:** Fangfang Yan, Akiko Suzuki, Chihiro Iwaya, Guangsheng Pei, Xian Chen, Hiroki Yoshioka, Meifang Yu, Lukas M. Simon, Junichi Iwata, Zhongming Zhao

## Abstract

The abnormal perturbation in gene regulation during palatogenesis may lead to cleft palate, a major congenital birth defect in humans and mice. However, a comprehensive multi-omic map of the developing secondary palate at single-cell resolution is lacking. In this study, we performed single-cell multiome sequencing and profiled chromatin accessibility and gene expression simultaneously within the same cells (n = 36,154) isolated from mouse secondary palate across embryonic days (E) 12.5, E13.5, E14.0, and E14.5. Application of optimal transport reconstructed five trajectories, representing continuous differentiation of multipotent cells into different subpopulations in later stages. By linking open chromatin signals to gene expression changes, we discovered a list of lineage-determining transcription factors, such as Shox2 for the anterior and Dlx1/2 for the posterior palatal mesenchymal trajectories. In conclusion, this study charted epigenetic and transcriptional dynamics during palatogenesis, which provides a valuable resource for the community and facilitate future research in cleft palate.

**Highlights:**

- The first report on building a single-cell multi-omics atlas with joint chromatin accessibility and gene expression measurements from the same cells during the development of mouse secondary palate.
- Application of optimal transport calculated fate probabilities to different terminal states and recovered continuous landscapes during mouse secondary palate development.
- By linking *cis*-regulatory DNA elements to target genes, we characterized a series of transcription factors governing the differentiation of cranial neural crest-derived multipotent cells to the anterior and posterior palatal mesenchymal trajectories, respectively.
- Transcription factors Shox2 and Dlx1/2 exhibited top regulatory roles for the anterior and posterior palatal mesenchymal trajectories, respectively, showing significant enrichment in both motif accessibility and gene expression.

## INTRODUCTION

The development of the secondary palate in both humans and mice is a dynamic, complex, and highly orchestrated three-dimensional process, involving vertical outgrowth, horizontal elevation, and fusion of palatal shelves (Bush and Jiang, 2012). Specifically, it begins around embryonic day (E) 12.5 in mice, when the palatal shelves initially arise from the lateral side of maxillary processes (Li et al., 2017). The palatal shelves grow vertically downward along with the tongue (E12.5-E13.5) and elevate to the horizontal position (E14.0). Following the elevation, the bilateral palatal shelves grow towards the midline and fuse into the intact palate (E14.5-E16.5) (Bush and Jiang, 2012; Li et al., 2017).

The whole process of palatogenesis is programmed by the precise control of gene expression by transcription factors (TFs) through binding to the accessible *cis*-regulatory DNA elements (CREs) of the target genes (Latchman, 1997). Abnormal perturbations may lead to cleft palate (CP), a major congenital birth defect in humans with a significant long-term impact on patients’ life quality (Dixon et al., 2011; Suzuki et al., 2016). Numerous efforts have aimed to uncover and annotate the CP-associated genes (Butali et al., 2019; Cai et al., 2017; Xu et al., 2021; Yan et al., 2020a), as well as the underlying regulatory mechanisms by integrating gene and microRNA expression data profiled from bulk tissue (Li et al., 2019a; Yan et al., 2020b; Yan et al., 2022). Beyond bulk tissue level, single-cell RNA-sequencing (scRNA-seq) technologies have been applied to study the palate and revealed large heterogeneity within the tissue. Recently, Han et al. mapped cell types and characterized lineage commitment by generating and analyzing time-series scRNA-seq data of the soft palate (Han et al., 2021). Another study defined the expression profiles of subpopulations in the fusing upper lip and primary palate at single-cell resolution (Li et al., 2019b).

While scRNA-seq reveals the transcriptional state differences between cells with high resolution, yet it provides little insight into the upstream regulations that drive such change (Chen et al., 2019). Single-cell epigenome assays, such as single-cell assay for transposase-accessible chromatin with sequencing (scATAC-seq), capture open chromatin signals and decipher the regulation status (Buenrostro et al., 2015). Computational approaches have been developed to integrate independent scRNA-seq and scATAC-seq datasets of the same tissue (Stuart et al., 2019). However, inferring “anchors” between datasets may not fully recapitulate the true molecular processes (Chen et al., 2019). More recently, single-cell multi-omics technologies have emerged as a powerful approach to accurately deciphering regulation status. They profile gene expression and chromatin accessibility simultaneously within the same cells. The epigenetic changes at the DNA level can be directly linked to transcriptomic changes at the RNA level to reveal the interplay between regulatory DNA sequences and the expression of target genes. Several multi-omics technologies, such as sci-CAR (Cao et al., 2018), Paired-seq (Zhu et al., 2019), SNARE-seq (Chen et al., 2019), and scCAT-seq (Liu et al., 2019), have been developed and applied to several tissue types, including human cerebral cortex (Trevino et al., 2021), human lung cancer (Liu et al., 2019), mouse kidney (Cao et al., 2018), and mouse embryos (Argelaguet et al., 2022). However, a comprehensive multi-omic map of gene expression and its regulation during the development of mouse secondary palate at single-cell resolution is still lacking.

In this study, we performed single-cell multiome sequencing (10x Multiome) and profiled the transcriptome and epigenome simultaneously within the same cells isolated from the developing mouse secondary palate spanning four critical developmental stages. A total of eight major cell types were identified, which were defined by canonical marker gene expression. By mapping open chromatin signals to gene expression changes, we discovered a list of cell-type specific regulators with enriched motif accessibility, as well as gene expression. We then focused on cranial neural crest (CNC)-derived multipotent cells, reconstructed five developmental trajectories, and uncovered a list of lineage-determining TFs that control the differentiation of each trajectory. This work represents the first report that building a single-cell multi-omic atlas of the developing mouse secondary palate. Insights into transcriptome and epigenome changes will increase our understanding of the underlying molecular processes and provides a valuable resource for the community.

## RESULTS

### Single-cell multiome assays dissect transcriptome and epigenome changes of the developing mouse secondary palate

The mouse secondary palate development mainly occurs between E12.5 and E14.5 (**Figure 1A**). To dissect gene regulation at the cellular level in the developing mouse secondary palate, we performed single-cell multiome sequencing using the 10x Chromium Single Cell Multiome platform and generated single-nucleus RNA-seq (snRNA-seq) and single-nucleus ATAC-seq (snATAC-seq) libraries from the same cells at E12.5 (n=2), E13.5 (n=3), E14.0 (n=2), and E14.5 (n=2) (**Figure 1A**). Jointly applying filters on both assays resulted in 36,154 cells with high-quality measurements across 36,155 genes and 123,807 accessible peaks representing potential CREs (**Figure S1A, Table S1**). The majority of cells had both high transcriptional start site (TSS) enrichment scores and a large number of fragments (**Figure S1B**). In addition, we observed nucleosome binding pattern (**Figure S1C, top**) and enrichment of chromatin accessibility around TSS compared to the flanking regions (**Figure S1C, bottom**). Together, these data suggested high snATAC-assay quality. Furthermore, biological replicates from the same developmental stage were highly correlated for both RNA (**Figure S1D, top,** coefficients = 1, *p*-values < 2.2×10^-16^) and ATAC measurements (**Figure S1D, bottom,** coefficients = 1, *p*-values < 2.2×10^-16^).

**Figure 1.**
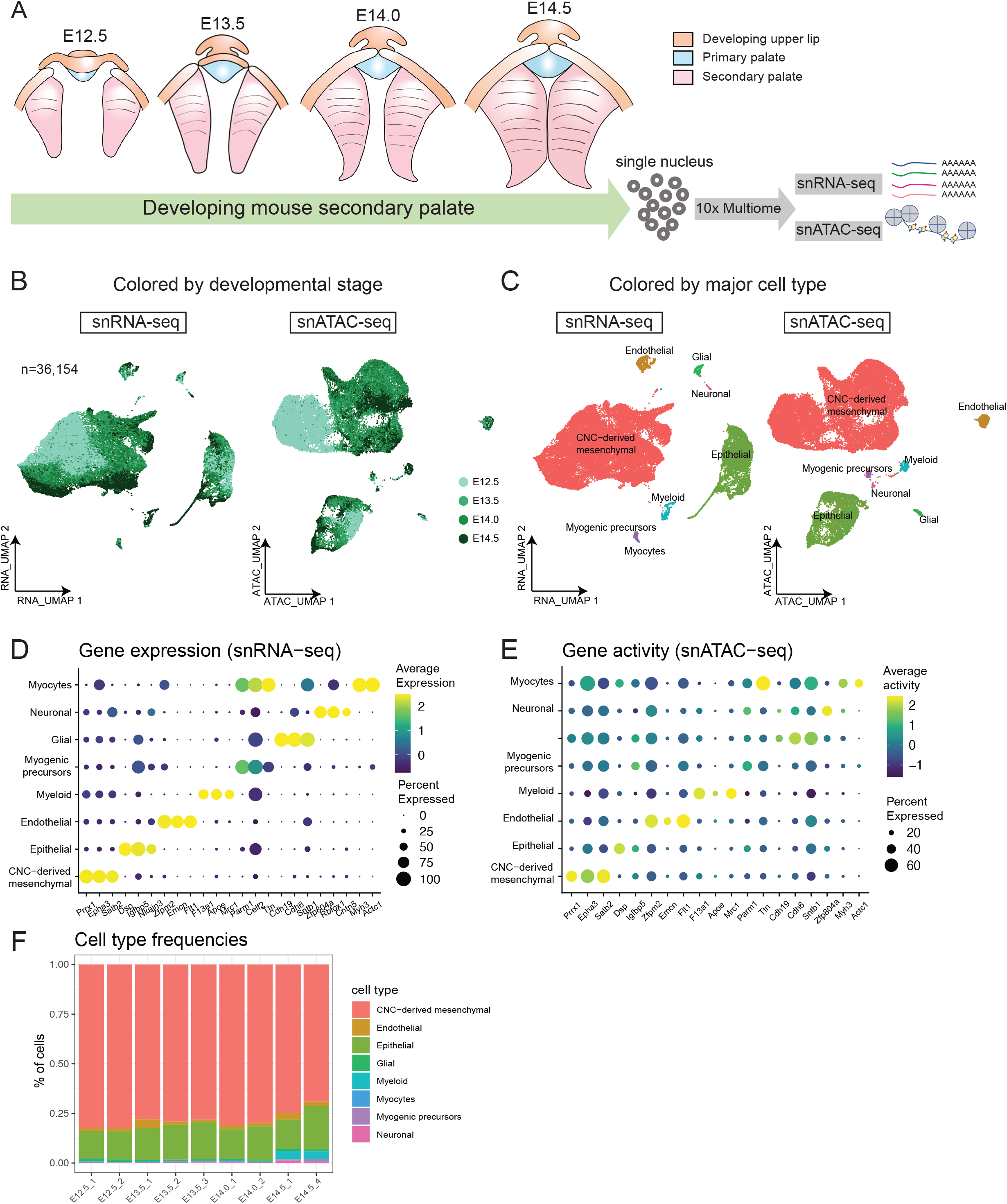
Single-cell multiome assays dissect transcriptome and epigenome changes of the developing mouse secondary palate. **(A)** Schematic plot depicts the development of mouse secondary palate (purple) spanning embryonic day (E) 12.5 (n=2), E13.5 (n=3), E14.0 (n=2), and E14.5 (n=2). The isolated nuclei are subjected to 10x Chromium Multiome sequencing to profile snRNA-seq and snATAC-seq simultaneously within the same cells. **(B)** Uniform manifold approximation and projection (UMAP) visualization of 36,154 cells based on RNA assay (left) or ATAC assay (right). Each dot represents one cell and is colored by the developmental stage. **(C)** Same UMAP visualization as panel B colored by annotated cell types. **(D, E)** Dot plot illustrates marker gene expression (x-axis) **(D)** and gene activity **(E)** (x-axis) across cell types (y-axis). Dot size is proportional to the percent of expressed cells. Colors indicate low (grey) to high (blue) expression. **(F)** Stacked bar plot shows cell type frequencies in each sample. Colors represent cell types.

Unsupervised dimension reductions based on gene expression (snRNA-seq) or chromatin accessibility (snATAC-seq) profiles revealed similar structures (STAR Methods). Within each distinct cluster, cells from E12.5 showed strong overlap with cells from later time points that occupied the borders of each cluster, suggesting an increase of complexity with palate development (**Figure 1B**). Individual cells of the biological replicates for each stage showed strong overlaps with each other (**Figure S1E**).

Next, we performed clustering analysis followed by cell type annotation. Compared to scRNA-seq, there is limited knowledge of cell-type specific open chromatin regions. Therefore, cell type annotation is more challenging in scATAC-seq data(Cusanovich et al., 2018). The frequently used computational approach involves cross-modality integration and label transfer from reference scRNA-seq data(Baek and Lee, 2020). Here, single-cell multi-omics technologies eliminates the need for inferring relationships *in silico*, which allows direct annotation of scATAC-seq-based clusters using scRNA-seq labels. We observed distinct clusters in both RNA and ATAC data, representing eight major cell types (**Figure 1C**). Each cell type was defined by canonical marker gene expressions, including CNC-derived mesenchymal cells (*Prrx1*, n=28,529, 78.91%), epithelial cells (*Krt14, Epcam*, n=5,866, 16.23%), endothelial (*Cdh5, Cldn5*, n=714, 1.97%), myeloid (*Lyz2*, n=397, 1.10%), glial cells (*Plp1, Sox10*, n=307, 0.85%), myogenic progenitors (*Myod1*, n=200, 0.55%), neuronal (*Tubb3, Stmn2*, n=113, 0.31%) and myocytes (*Myf5*, n=28, 0.08%) (**Figure 1D, Figure S2A**). We then quantified the gene activity score, a metric defined by aggregating accessible chromatin regions intersecting the gene body and promoters in snATAC-seq data. The abovementioned markers, which showed cell type-specific expression, also exhibited similar patterns of chromatin accessibility in corresponding clusters (**Figure 1E**). We then examined cell type frequencies and found that each cluster included cells from all time points without strong bias for earlier or later stages (**Figure 1F**).

To further corroborate our inferred cell type identities, we downloaded a recently published scRNA-seq dataset of the developing mouse soft palate (Han et al., 2021). Following normalization and dimension reduction, we projected the soft palate data into our scRNA-seq manifold using canonical correlation analysis (Stuart et al., 2019) (STAR Methods) and observed high agreement between cell type annotations (**Figure S2B-C**).

### Direct linking of open chromatin signals to gene expression discovers cell typespecific regulators

Paired RNA and ATAC measurements within the same cells reveal both the transcriptional state and the upstream DNA regulatory element activities, which allow direct mapping of epigenetic gene regulation to gene expression. We aimed to identify TFs with both enriched accessibility and expression profiles for a specific cell type, representing putative regulators for that cell type. Towards this end, we first conducted differential gene expression analysis between cell types and identified 6,573 differentially expressed genes (DEGs) (adjusted *p*-value < 0.05, log Fold change > 0.1, min.pct > 0.1). To find regulatory elements for each DEG, we conducted CRE-gene linkage analysis by calculating the Pearson correlation coefficient (PCC) between CRE accessibility and gene expression while accounting for CRE size and fragment count (STAR Methods). Positively linked CRE-gene pairs may represent enhancer-gene interactions. A total of 15,018 pairs were identified, including 12,596 CREs significantly linked to 3,787 cell type-specific genes (**Figure 2A**, score > 0, adjusted *p*-value < 0.05). Each cell-type specific gene was linked to a median of three CREs (min=1, max=28, mean=3.966). For example, in CNC-derived mesenchymal cells, *Twist1* expression was linked to 15 CREs. Of these locus chr12:33957146-33958061 was the most significant one (score = 0.315, adjusted *p*-value = 1.70×10^-8^). Both the expression level of *Twist1* and accessibility of the CRE chr12:33957146-33958061 were increased in CNC-derived mesenchymal cells compared to all other cell types (**Figure 2B**). Genome browser visualization of the *Twist1* locus revealed that chr12:33957146-33958061 partially overlaps the transcription start site of *Twist1*. Therefore, chr12:33957146-33958061 most likely acts as an enhancer that upregulates the expression of *Twist1* in CNC-derived mesenchymal cells (**Figure 2B**). Another exemplary CRE-gene pair was *Tie1* and locus chr4:118489480-118490171, showing significant enrichment in endothelial cells (score = 0.369, adjusted *p*-value = 3.88×10^-14^, **Figure S3**).

**Figure 2.**
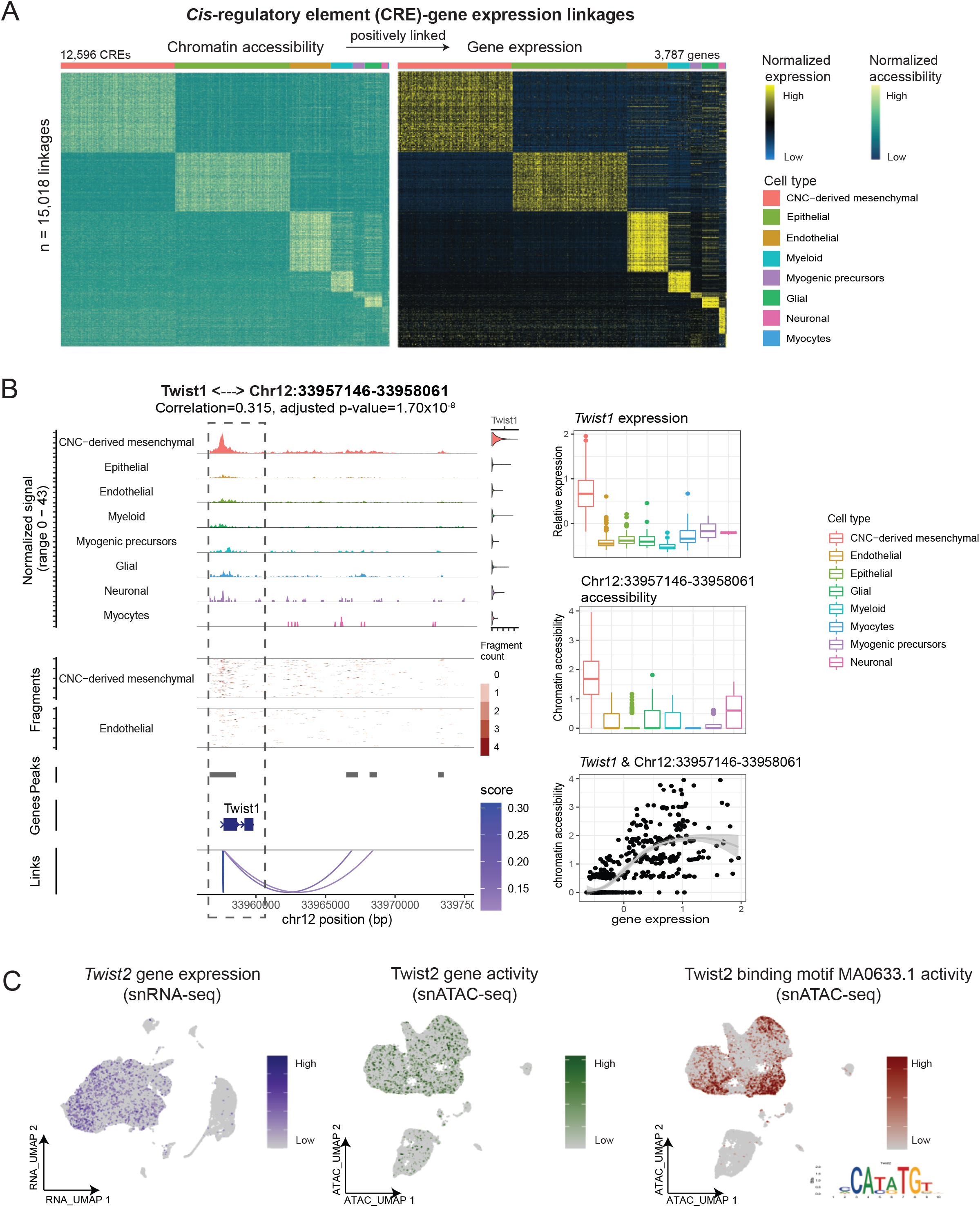
Direct linking of open chromatin signals to gene expression discovers cell type-specific regulators of secondary palate formation. **(A)** Heatmap shows normalized chromatin accessibility (left) and gene expression (right) of 15,018 significantly linked cis-regulatory elements (CREs)-gene pairs. Each row represents a positively linked pair of CREs and genes. Bar on the top represents major cell types. **(B) Left**: Genome Browser visualization of aggregated chromatic accessibility at the chr12: 33.96-33.97 (Mb) locus for each major cell type, coupled with *Twist1* gene expression. Arcs at the bottom denote positively linked *CRE-Twist1* pairs. The linkage between *Twist1* and chr12:33957146-33958061 is highlighted with blue dotted box. **Right**: Boxplots show the expression level of *Twist1* (top) and accessibility of the linked CRE chr12:33957146-33958061 (middle) across cell types. Scatter plot shows the significant correlation between *Twist1* expression and chr12:33957146-33958061 accessibility (bottom). **(C)** UMAP visualizations illustrate the multimodal profiling of *Twist2* including gene expression (left), gene activity (middle), and Twist2 binding motif MA0633.1 activity (right). The position weight matrix of the MA0633.1 motif is embedded in the bottom right corner.

To nominate TFs that control each major cell type, we identified enriched motifs of these CREs. The following criteria were applied to define cell-type specific regulatory TFs: (1) TF expression was enriched at the RNA level, and (2) TF binding motif accessibility was enriched in the ATAC measurements. In total, we discovered 81 putative regulators for the eight major cell types (adjusted RNA *p*-value < 0.05 & adjusted motif *p*-value < 0.05, **Table 1**). For example, both the RNA expression of *Twist2* (*RNA.p-value* < 2.22×10^-16^) and chromatin accessibility of the *Twist2* binding motif MA0633.1 (*motif.p-value* = 4.91×10^-18^) were enriched in CNC-derived mesenchymal cells (**Figure 2C**). In endothelial cells, we found enrichment of both the expression of the *Foxo1* (*RNA.p-value* < 2.22×10^-16^) and the accessibility of the its binding motif MA0480.1 (*motif.p-value* = 4.91×10^-18^).

**Table 1.** Summary of top five putative regulators for each major cell type.

| Major cell type | Gene symbol | RNA.p-value | Motif | Motif.p-value |
| --- | --- | --- | --- | --- |
| CNC-derived mesenchymal | *Twist2* | 3.00E-241 | MA0633.1 | 0 |
| CNC-derived mesenchymal | *Lhx8* | 0 | MA0705.1 | 2.01E-61 |
| CNC-derived mesenchymal | *Foxf1* | 2.97E-192 | MA1606.1 | 5.41E-133 |
| CNC-derived mesenchymal | *Shox2* | 6.76E-211 | MA0720.1 | 1.72E-73 |
| CNC-derived mesenchymal | *Tcf3* | 2.01E-05 | MA0092.1 | 2.03E-257 |
| Epithelial | *Sox2* | 0 | MA0142.1 | 2.94E-88 |
| Epithelial | *Sox6* | 2.16E-225 | MA0515.1 | 2.05E-197 |
| Epithelial | *Smad3* | 1.38E-144 | MA1622.1 | 8.26E-215 |
| Epithelial | *Klf12* | 1.00E-09 | MA0742.1 | 0 |
| Epithelial | *Runx1* | 2.58E-233 | MA0002.2 | 1.30E-36 |
| Endothelial | *Foxo1* | 8.90E-145 | MA0480.1 | 4.91E-18 |
| Endothelial | *Nr2f6* | 7.22E-07 | MA0677.1 | 3.08E-113 |
| Endothelial | *Sox17* | 0 | MA0078.1 | 9.56E-30 |
| Endothelial | *Nr5a2* | 7.58E-145 | MA0505.1 | 5.63E-37 |
| Endothelial | *Esrrg* | 1.69E-15 | MA0643.1 | 1.39E-57 |
| Myeloid | *Runx1* | 6.16E-104 | MA0002.2 | 1.24E-08 |
| Myeloid | *Nfe2l2* | 1.67E-65 | MA0150.2 | 5.29E-29 |
| Myeloid | *Arnt* | 4.92E-4 | MA0004.1 | 9.49E-21 |
| Myeloid | *Bach1* | 7.34E-3 | MA0591.1 | 3.27E-23 |
| Myeloid | *Nr4a2* | 1.72E-3 | MA0160.1 | 5.49E-22 |
| Myogenic precursors | *Tcf12* | 2.09E-06 | MA0521.1 | 1.73E-115 |
| Myogenic precursors | *Tcf21* | 3.08E-68 | MA0832.1 | 5.97E-105 |
| Myogenic precursors | *Plagl1* | 2.07E-16 | MA1615.1 | 1.68E-3 |
| Myogenic precursors | *Arx* | 3.20E-63 | MA0874.1 | 7.88E-06 |
| Myogenic precursors | *Nobox* | 3.38E-4 | MA0125.1 | 4.40E-06 |
| Glial | *Sox5* | 8.83E-146 | MA0087.1 | 1.65E-4 |
| Glial | *Nr4a2* | 5.35E-48 | MA0160.1 | 2.39E-42 |
| Glial | *Creb5* | 1.91E-63 | MA0840.1 | 3.30E-22 |
| Glial | *Dlx1* | 1.64E-138 | MA0879.1 | 1.23E-4 |
| Glial | *Rxra* | 3.49E-4 | MA0065.2 | 3.97E-52 |
| Neuronal | *Arid3b* | 1.37E-06 | MA0601.1 | 3.29E-05 |
| Neuronal | *Hmx2* | 5.68E-177 | MA0897.1 | 8.05E-3 |
| Neuronal | *Zic1* | 1.03E-06 | MA1628.1 | 8.96E-3 |

### CNC-derived mesenchymal subpopulations correspond to in vivo anatomical locations

To understand the heterogeneity within CNC-derived mesenchymal cells, we isolated this cluster and conducted an independent analysis, including normalization, clustering, and dimension reduction (**Figure 3A, left panel**). In the dimension-reduced data manifold, we observed a continuous progression from E12.5 to E14.5 (**Figure 3A, S4A**). The cells from the early stage (E12.5), which resided near the middle of the lowdimensional space, were more homogeneous compared to cells from later stages. Cells from the late stage (E14.5) represented differentiated cells residing at the edge of the embedding. Cell subtype annotation was conducted through extensive manual curation of marker genes. We identified seven subtypes characterized by specific gene expression signatures (**Figure 3A, right panels**). For example, anterior palatal mesenchymal cells exhibited high expression of short stature homeobox 2 (*Shox2*) (Li and Ding, 2007) and ALX Homeobox 1 (*Alx1*) (**Figure S4B**). Chondrogenic cells were characterized by high expression of *Sox9* and *Col12a1* (Izu et al., 2011) while osteoblasts had high expression of *Runx2* and *Sp7* (Yuan et al., 2020). Dental mesenchymal cells exhibited high expression of *Dlx2, Sostdc1* (Munne et al., 2009), and *Tfap2b* (Woodruff et al., 2021). Posterior palatal mesenchymal cells had high expression of *Tbx22* and *Wnt16* (Han et al., 2021). A list of progenitor-related genes was highly expressed in cluster 5, such as *Dach1, Lmo4, Hmgb2, Hmgb3*, and *Runx1t1* (adjusted *p*-values < 2.2×10^-16^). Gene Set Enrichment Analysis (GSEA) suggested these genes were significantly associated with regulation of stem cell proliferation (GO: 0072091, FDR = 2.58×10^-3^, enrichment ratio = 92.87; FDR: false discovery rate) and enriched in neural progenitor cells (FDR = 2.68×10^-3^, enrichment ratio = 13.97) (**Figure S4C**). Thus, cluster 5 was annotated as CNC-derived multipotent cells.

**Figure 3.**
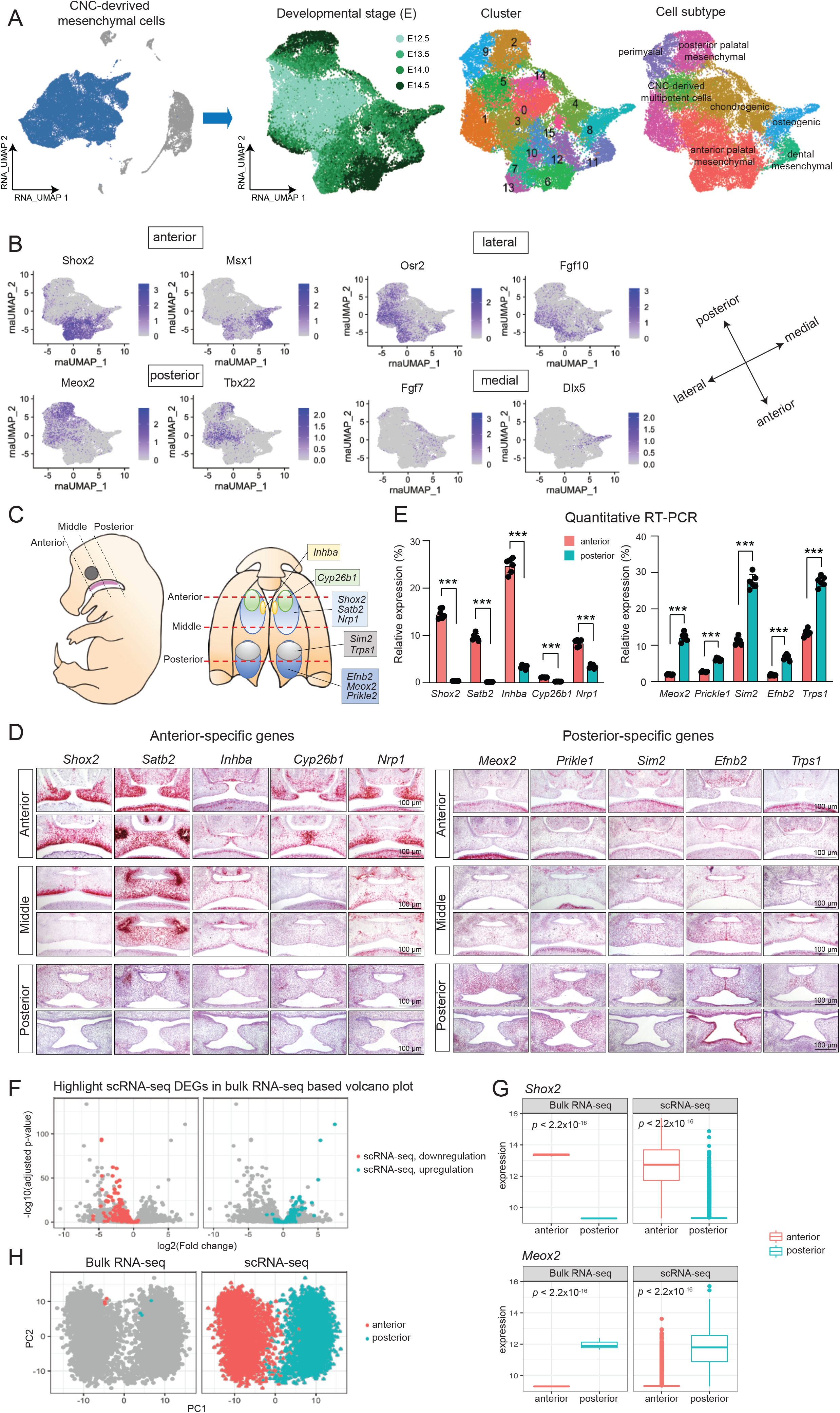
CNC-derived mesenchymal subpopulations correspond to *in vivo* anatomical locations. **(A) Left:** UMAP visualization of the whole dataset with CNC-derived mesenchymal cells highlighted in blue. **Right**: Independent UMAP visualizations of CNC-derived mesenchymal cells colored by developmental stage, cluster, and cell subtype, respectively. **(B)** Feature plot shows the expression of representative genes in the anterior, posterior, lateral, and medial, respectively. **(C)** Anatomical mouse embryo images outline gene expression patterns by RNAscope *in situ* hybridization. **(D)** Microscope images (100 μm) show expression patterns for each gene by *in situ* hybridization. **(E)** Bar plot shows quantitative RT-PCR of genes in anterior (red bars) and posterior (blue bars) regions. ***p<0.001. **(F)** Volcano plot shows the log2 Fold Change and negative log10 of adjusted *p*-values for each gene in the bulk RNA-seq dataset and colored by significance in the single cell RNA-seq dataset. **(G)** Boxplot shows the expression of two representative DEGs, *Shox2* and *Meox2*, at the bulk level (left) and single-cell level (right). **(H)** Scatter plot shows the projection of bulk RNA-seq data into the PCA space defined by scRNA-seq data. Color represents anterior (red) or posterior (blue). As expected, bulk samples from different locations clustered with corresponding scRNA-seq samples.

Of note, gene expression patterns aligned with *in vivo* anatomical locations (**Figure 3B**). For example, *Shox2* and *Msx1* were specifically expressed in anterior regions while *Meox2* and *Tbx22* expression were restricted to posterior regions (Smith et al., 2013). *Osr2* and *Fgf10* exhibited expression in lateral regions while *Fgf7* and *Dlx5* were expressed in medial regions (Lan et al., 2004). Even though expression patterns of *Shox2* and *Meox2* have been well studied, the majority of region-specific genes we found in this study have not been well characterized in the developing palate (**Table 2**).

**Table 2.**
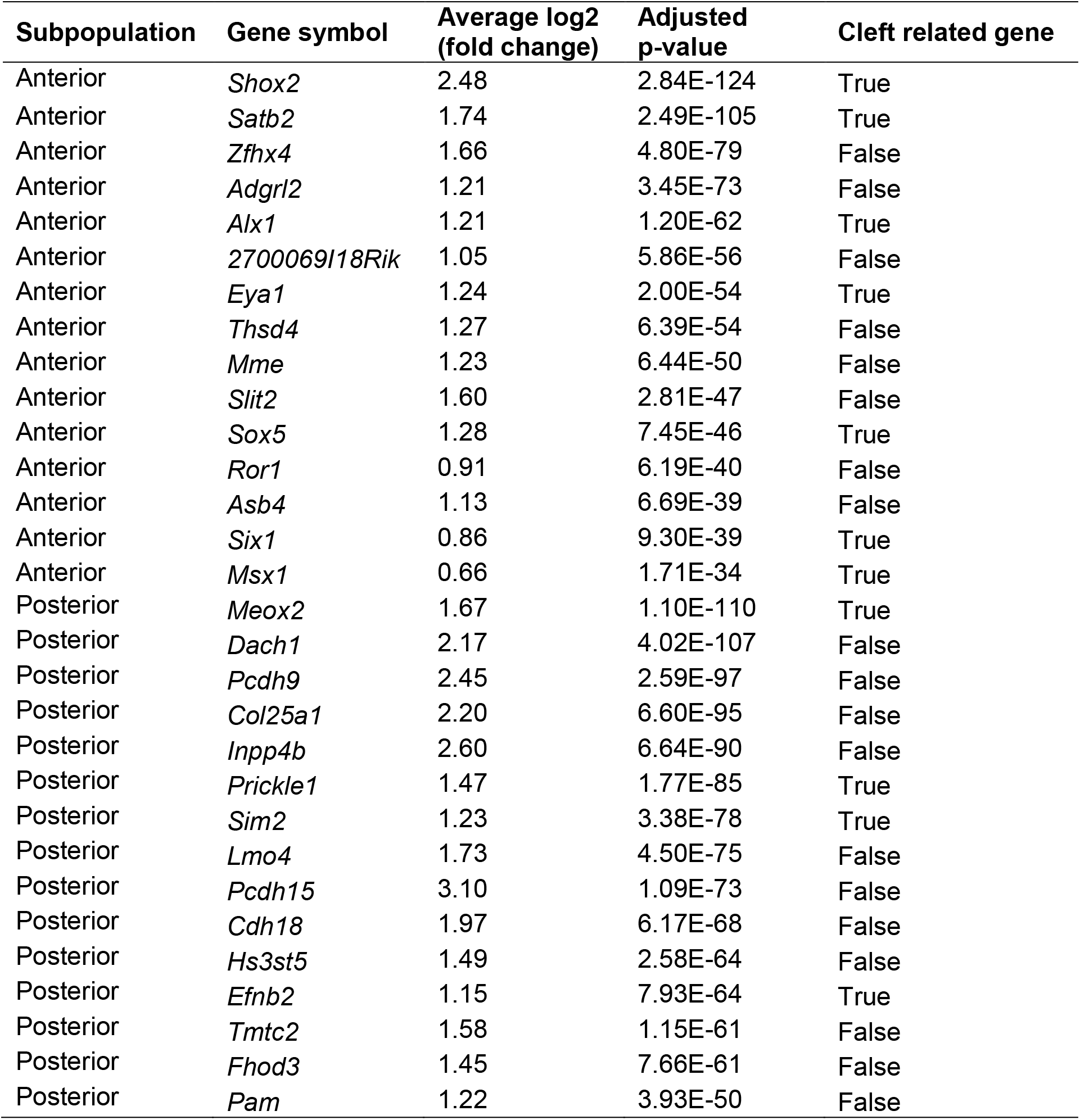
Summary of marker genes in anterior and posterior subpopulations.

To validate the *in vivo* identities of subpopulations, we conducted RNAscope *in situ* hybridization. The top five subpopulation-specific genes were selected, that is *Shox2, Satb2, Inhba, Cyp26b1*, and *Nrp1* for the anterior subpopulation, *Meox2, Prickle1, Sim2, Efnb2*, and *Trps1* for the posterior subpopulation (**Figure 3C, D**). It was found that *Shox2, Satb2*, and *Nrp1* were expressed mainly in the anterior to the middle region. The expression of *Cyp26b1* was restricted to the anterior region and *Inhba* was restricted to beneath the epithelial layer in the anterior region. In addition, *Meox2, Prikle1*, and *Efnb2* were expressed in the entire posterior region. *Sim2* and *Trps1* were expressed only in the anterior half of the posterior region of the developing secondary palate. These gene expression patterns were also validated with quantitative reversetranscription polymerase chain reaction (qRT-PCR) (**Figure 3E**).

For further validation, we performed bulk RNA sequencing using anterior (n=3) and posterior regions (n=3) of the developing secondary palate at E14.0. We observed consistent expression patterns in bulk RNA-seq and scRNA-seq data. The majority of upregulated genes (75/86, 87.21%) in the scRNA-seq anterior cluster were found to also be upregulated in the bulk RNA-seq anterior tissue (**Figure 3F**). Fisher’s Exact test revealed significant enrichment between scRNA-seq DEGs and bulk RNA-seq DEGs (odds ratio = 1354.47, *p*-value < 2.2×10^-16^). For example, *Shox2* was significantly higher expressed in the anterior (bulk RNA-seq adjusted *p*-value = 9.12×10^-49^, scRNA-seq adjusted *p*-value < 2.20×10^-16^), while *Meox2* exhibited higher expression in the posterior region (bulk RNA-seq adjusted *p*-value = 1.69×10^-94^, scRNA-seq *p*-value < 2.20×10^-16^, **Figure 3G**). In addition, we conducted principal component analysis (PCA) using the scRNA-seq DEGs and projected the bulk RNA-seq data into this PC space. As expected, bulk samples from different locations clustered with corresponding scRNA-seq transcriptomes (**Figure 3H**). Overall, these results validated our subtype annotations of CNC-derived mesenchymal cells.

### Reconstruction of CNC-derived mesenchymal trajectories by optimal transport reveals lineage-determining transcription factors

As chondrocytes originate from the pterygoid plate anlage and are not considered as part of the secondary palate (Han et al., 2021), we excluded them from downstream analysis (**Figure 4A**). Single-cell data from discrete time points can be considered “snapshots” of the underlying continuous developmental process (Luecken and Theis, 2019). To connect static “snapshots” into a “movie” and computationally reconstruct the molecular dynamics during the differentiation of CNC-derived mesenchymal cells, we applied Wadding-Optimal Transport (WOT) (Lange et al., 2022; Schiebinger et al., 2019), an algorithm designed for trajectory analysis of time series scRNA-seq data. WOT connects adjacent time points by finding the most probable cell transition paths using the mathematical theory of optimal transport (Kantorovitch, 1958; Monge, 1781). Simulated random walks based on the WOT-derived cell transition matrix showed that most trajectories started from CNC-derived multipotent cells (black dots) and terminated at various subpopulations at late stages (yellow dots) (**Figure 4B, left panel**). We then quantified the terminal state likelihood of each cell and defined those with high likelihoods as terminal state cells (**Figure 4B, right panel**). Next, we computed the probabilities that an early cell would transition towards any terminal state cells. Overall, we discovered five trajectories, representing the continuous differentiation of multipotent cells into (1) anterior palatal mesenchymal cells, (2) posterior palatal mesenchymal cells, (3) dental mesenchymal cells, (4) osteoblasts, and (5) perimysial cells (**Figure 4C, Figure S5**). We also calculated diffusion pseudotime (Haghverdi et al., 2016) and found consistent results (STAR Methods) (**Figure S6A**).

**Figure 4.**
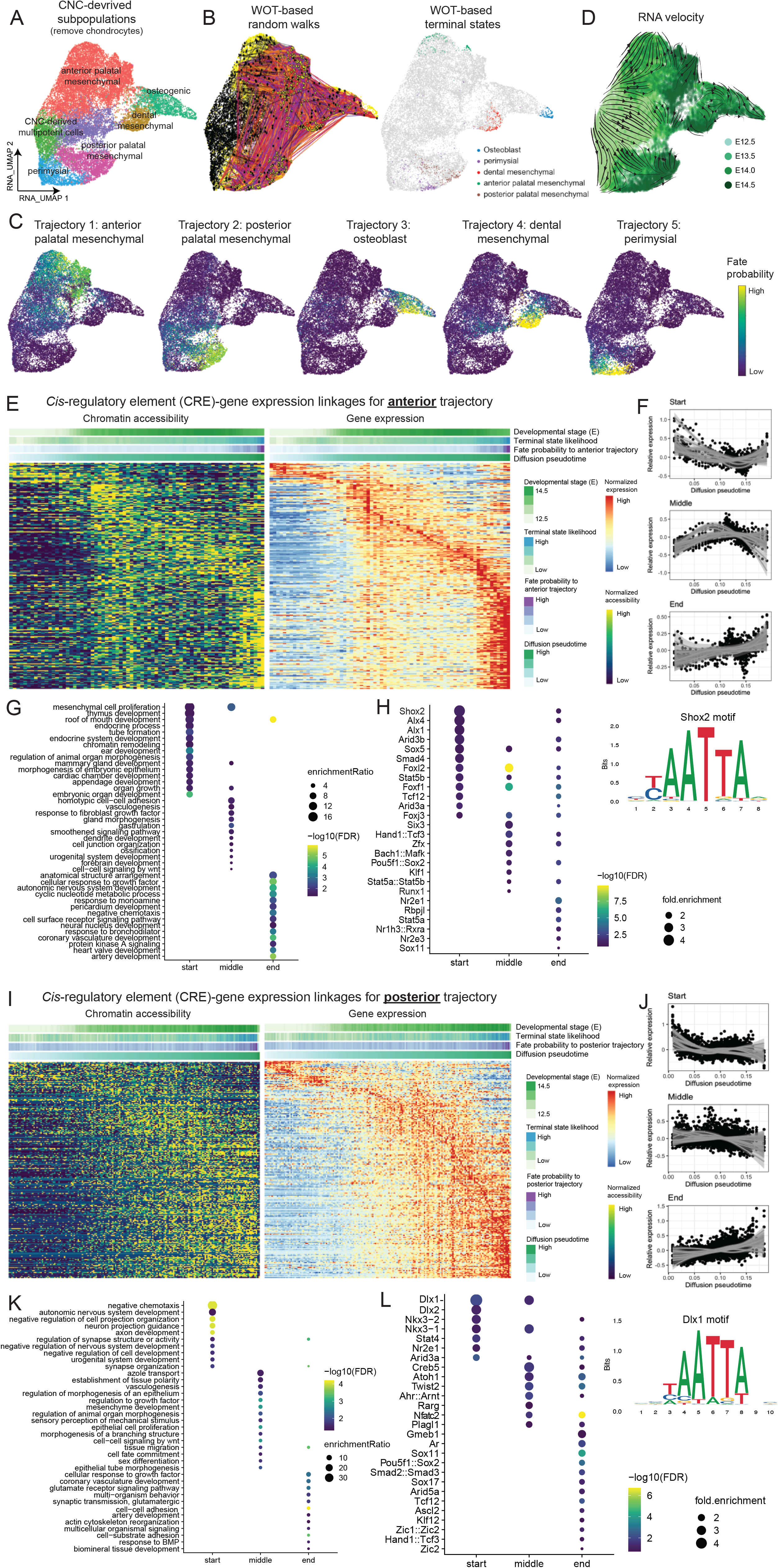
Reconstruction of CNC-derived mesenchymal trajectories by Optimal Transport reveals lineage-determining transcription factors. **(A)** UMAP visualization of CNC-derived mesenchymal subpopulations with chondrocytes being removed. **(B)** UMAP visualization colored by (left) WOT-based random walks and (right) terminal states. The random walks are based on the WOT-derived cell-cell transition matrix. Black dots represent the start points of the trajectory while yellow dots represent endpoints. **(C)** UMAP visualizations show fate probabilities to each trajectory. Cells are colored by probabilities (Yellow: high, Dark brown: low). **(D)** UMAP visualization with streamlines and arrows shows RNA velocity-derived information. Each point represents one cell and is colored by the developmental stage. Streamlines and arrows represent the future directions for each cell. **(E)** Heatmap shows paired chromatin accessibility and gene expression (rows) for the anterior trajectory (columns). Each row represents a putative pair of genes and linked regulatory elements. Bars on the top represent diffusion pseudotime, fate probabilities to the anterior palatal mesenchymal trajectory, terminal state likelihood, and developmental stage. Columns are ordered by diffusion pseudotime. **(F)** Scatter plot with fitted lines shows the expression pattern for each group of driver genes along the trajectory. **(G)** Dot plot shows enriched pathways (y-axis) for each driver gene group (x-axis). Dot is scaled by enrichment ratio and colored by significance. **(H)** Dot plot shows enriched motifs (y-axis) at different stages of the anterior trajectory (x-axis). Dot is scaled by motif enrichment ratio and colored by significance. The position weight matrix of one representative motif Shox2 is included. **(I-L)** Similar visualizations to panels E-H for the posterior palatal mesenchymal trajectory.

To validate inferred trajectories, we calculated RNA velocity, a measure to predict future states of individual cells based on the stratification of spliced and unspliced mRNAs (La Manno et al., 2018). As shown in **Figure 4D**, the directed dynamic information based on RNA velocity was consistent with inferred trajectories. The RNA velocities pointed away from the cells at E12.5 in the middle of the embedding towards later time points at the border of the embedding (**Figure 4D**). To more granularly resolve these velocity predictions, we applied CellRank, which infers initial and terminal states probabilities for each cell based on RNA velocity (Lange et al., 2022). Consistent with WOT-derived trajectories, CellRank found high initial state probabilities in CNC-derived multipotent cells and high terminal state probabilities in the late-stage subpopulations (**Figure S6B**). To eliminate potential bias caused by developmental stages, we repeated the analysis restricted to cells from E12.5 only and observed consistent results (**Figure S6C**).

We next investigated both gene expression and regulation dynamics of each trajectory. We first focused on the anterior palatal mesenchymal trajectory (**Figure 4EH**). Cells with large diffusion pseudotime values tended to be derived from late stages with high terminal state likelihood (**Figure S7A**). To pinpoint the driver genes for this trajectory, we conducted an association test between the expression level of each gene and fate probability. Those with significant positive correlations were defined as driver genes. We identified a total of 556 driver genes (correlation > 0.05 and adjusted *p*-value < 0.05). The top hits included *Shox2* (correlation = 0.468, adjusted *p*-value < 2.2×10^-16^), *Foxd2os* (correlation = 0.391, adjusted *p*-value < 2.2×10^-16^), and *Foxd2* (correlation = 0.333, adjusted *p*-value < 2.2×10^-16^, **Figure S7B**). To further investigate when and how these driver genes were regulated along the trajectory, we extracted 7,240 cells with high probabilities to differentiate towards the anterior trajectory (fate probability > 75% quantile) and ordered them by diffusion pseudotime. We then performed CRE-gene linkage analysis as described above and connected expression trajectories with chromatin accessibility dynamics. Out of 984 CRE-gene linkages, 428 (43.49%) were significantly linked (correlation > 0 and adjusted *p*-value < 0.05). We observed consistent gene expression and chromatin accessibility dynamics along the anterior trajectory (**Figure 4E**).

Using k-means clustering, these driver genes were divided into three groups, showing high expression at the start, middle, and end stages of the anterior trajectory, respectively (**Figure 4F**). The genes that were upregulated at the start of the anterior trajectory were enriched in mesenchymal cell proliferation (enrichment ratio = 16.71, adjusted *p*-value = 0.036), roof of mouth development (enrichment ratio = 12.73, adjusted *p*-value = 0.022), and chromatin remodeling (enrichment ratio = 9.90, adjusted *p*-value = 0.035, **Figure 4G**). The genes upregulated at the middle of the trajectory were enriched in mesenchymal cell proliferation (enrichment ratio = 11.14, adjusted *p*-value = 2.28×10^-3^) and response to fibroblast growth factor (enrichment ratio = 5.74, adjusted *p*-value = 0.027) while the genes upregulated at the end of the trajectory were associated with the roof of mouth development (enrichment ratio = 7.49, adjusted *p*-value = 1.04×10^-6^) and anatomical structure arrangement (enrichment ratio = 10.49, adjusted *p*-value = 4.58×10^-3^). By applying the motif enrichment test and setting multiple criteria, we characterized a list of lineage-determining TFs that control the anterior trajectory, such as Shox2 at the early stage (motif adjusted *p*-value = 6.09×10^-3^, motif fold change = 4.21, gene adjusted *p*-value < 2.2×10^-16^, gene fate correlation = 0.47), Foxl2 at the middle stage, and Nr2e1 at the late stage of the trajectory (**Figure 4H**).

We then applied analogous approach to the posterior palatal mesenchymal trajectory and identified 586 driver genes. Among them, 216 genes were significantly regulated by 353 CREs (**Figure 4I**), including *Col25a1* (correlation = 0.518, adjusted *p*-value < 2.2×10^-16^), *Meox2* (correlation = 0.496, adjusted *p*-value < 2.2×10^-16^), and *Inpp4b* (correlation = 0.400, adjusted *p*-value < 2.2×10^-16^) (**Figure S7C**). Pathway enrichment analysis of these genes revealed the involvement of neuron-related pathways in the early stage, mesenchymal development (enrichment ratio = 5.42, adjusted *p*-value = 1.37×10^-4^), and tissue migration (enrichment ratio = 3.57, adjusted *p*-value = 4.09×10^-4^) in the intermediate and late stages of the trajectory (**Figure 4J, K**). Transcription factor Dlx1 showed early regulatory roles for the posterior trajectory (motif adjusted *p*-value = 8.53×10^-3^, motif fold change = 4.78, gene *p*-value = 2.18×10^-4^, gene fate correlation = 0.02, **Figure 4L**).

To further validate these predictions, we first examined the odds ratio distribution of fate probabilities towards anterior versus posterior trajectories. As expected, *Shox2-* positive cells had high probabilities to differentiate towards the anterior palatal mesenchymal trajectory (**Figure 5A, left panel**) while *Meox2*-positive cells had high fate probabilities towards the posterior trajectory (**Figure 5B, left panel**). Those terminally differentiated cells at E14.5 emerged from the multipotent cells at the early stage (E12.5) (**Figure 5AB, right panels**). In addition, we observed a significant negative correlation of fate probabilities between the anterior and posterior trajectories (PCC Rho = −0.427, *p*-value < 2.2×10^-16^, **Figure 5C**). For example, the top driver gene for anterior trajectory *Shox2* was negatively correlated with the posterior trajectory (correlation = −0.333, adjusted *p*-value < 2.2×10^-16^). *Meox2*, a top driver gene for the posterior trajectory, was negatively correlated with the anterior trajectory (correlation = −0.288, adjusted *p*-value < 2.2×10^-16^). The osteoblast and dental mesenchymal trajectory shared a list of driver genes, exhibiting high fate probabilities of both trajectories, such as *Runx2* and *Zfpm2* (**Figure 5D**). Collectively, these data validated the inferred trajectories and driver genes.

**Figure 5.**
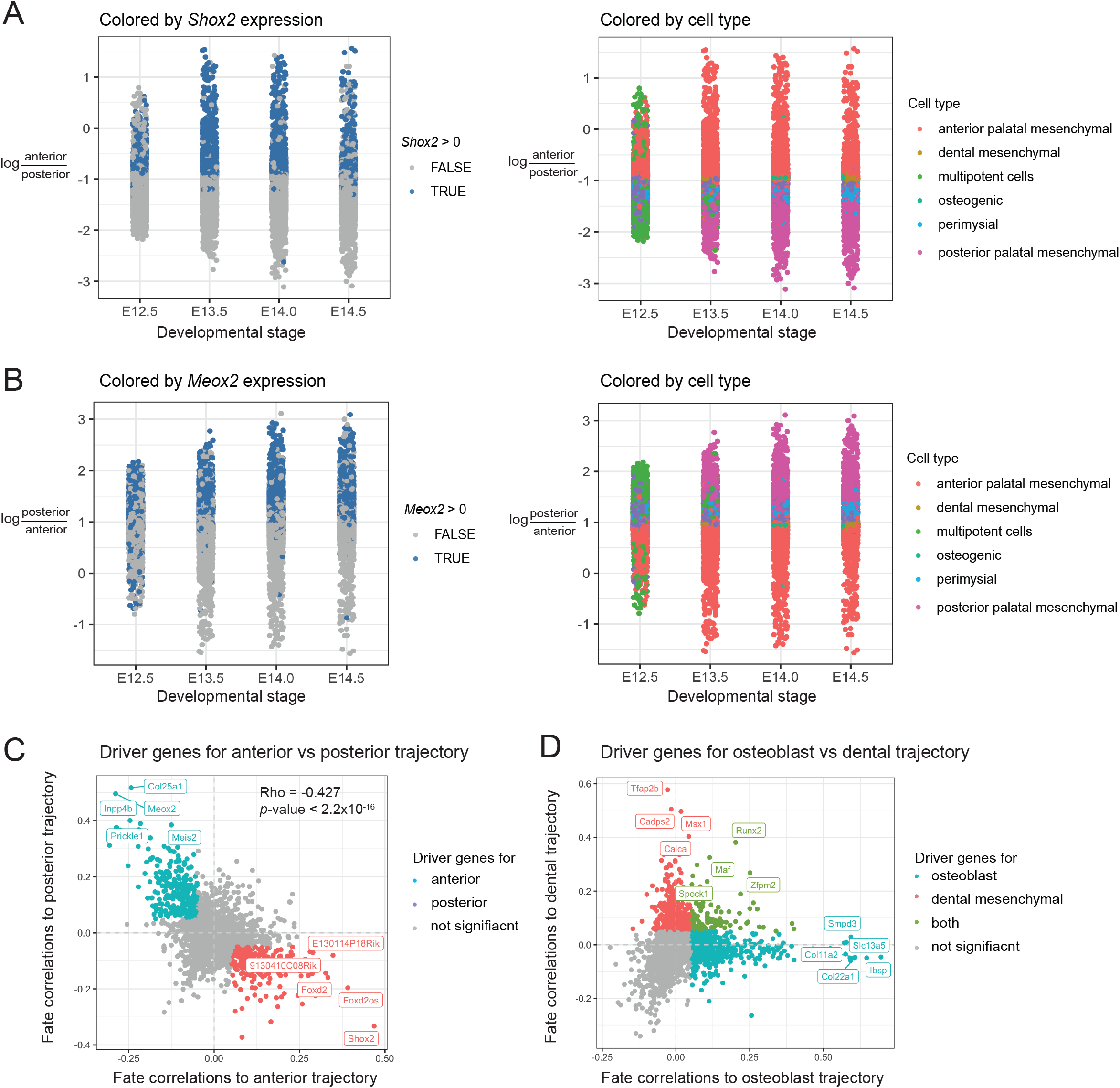
Driver genes for the anterior trajectory show negative fate correlations to the posterior trajectory, which validate the inferences. **(A)** Dot plot shows the distribution of log odds of fate probabilities to anterior versus posterior trajectories (y-axis) in each cell across the developmental stage (x-axis). Each dot represents one cell and is colored by *Shox2* expression (left) or cell types (right). Cells with positive expression of *Shox2* also exhibited high fate probabilities to anterior compared to the posterior trajectory. (**B**) Similar visualization to panel A for log odds of fate probabilities to posterior versus anterior trajectories. Each dot represents one cell and is colored by *Meox2* expression (left) or cell types (right). Cells with positive expression of *Meox2* also exhibited high fate probabilities to anterior compared to the posterior trajectory. **(C)** Scatter plot shows fate correlations to anterior (x-axis) and posterior (y-axis) trajectories of each gene. Defined driver genes for each trajectory are highlighted in the plot. **(D)** Scatter plot shows fate correlations to osteoblast (x-axis) and dental mesenchymal (y-axis) trajectories of each gene. Genes in green have similar fate correlations to both trajectories.

### Epithelium is differentiated into oral, nasal, and dental epithelium

We then focused on epithelial cells and annotated the subtypes using markers from previous publications (**Figure 6A, left panel**). Clusters 0 and 1 were labeled as undifferentiated epithelium with positive expression of *Trp63* and negative expression of Keratins. Clusters 2 and 7 were characterized by high expression of dental epithelium markers, such as *Fst, Fgf9*, and *Jag2* (Mitsiadis et al., 2010). Clusters 4 and 8, annotated as nasal epithelium, exhibited co-expression of cilia- and flagella-associated protein genes (*Cfap299* and *Cfap47*)(*Ahn* et al., 2021) and Keratins (*Krt15*). Clusters 3 and 11 had high expression of *Krt19*, which is a periderm-specific gene (Richardson et al., 2014) (**Figure 6A, right panels**). Cluster 6 specifically expressed *Shh* and was annotated as the oral epithelium of the palatal rugae (Sagai et al., 2017).

**Figure 6.**
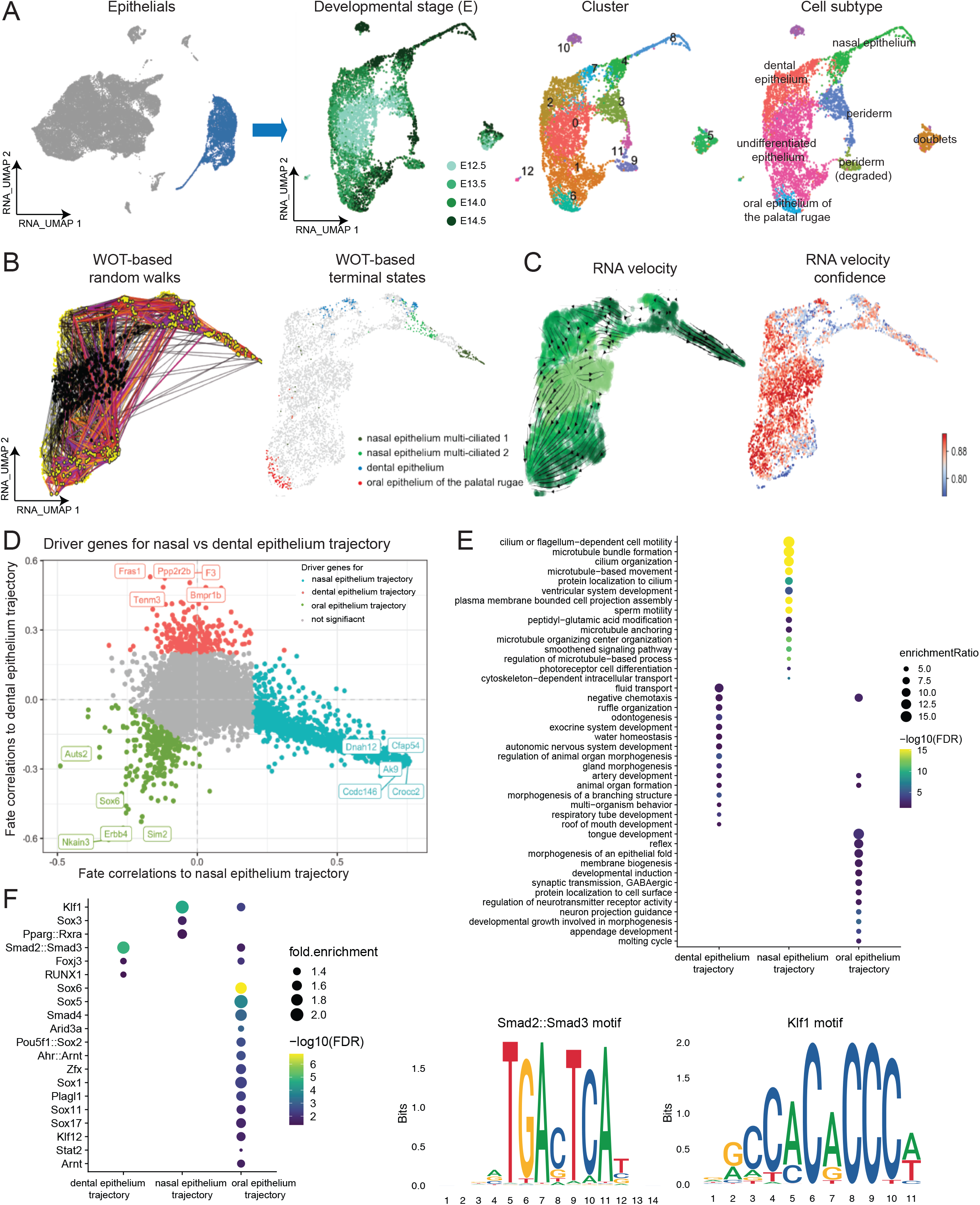
Epithelium is differentiated into the oral, nasal, and dental epithelium. **(A)** Left: UMAP visualization of the whole dataset with epithelium highlighted in blue. Right: Independent UMAP visualizations of epithelium colored by developmental stage, cluster, and cell subtype, respectively. **(B)** UMAP visualization colored by (left) WOT-based random walks and (right) terminal states. The random walks were based on the cell-cell transition matrix. Black dots represent start points and yellow dots represent endpoints. **(C)** UMAP visualization with streamlines and arrows showing RNA velocity-derived information. Each point represents one cell and is colored by developmental stage (left) or velocity confidence (right). Streamlines and arrows represent the future directions for each cell. **(D)** Scatter plot shows fate correlations to the dental epithelium (x-axis) and nasal epithelium (y-axis) trajectories of each gene. Defined driver genes for each trajectory are highlighted in the plot. Blue: dental epithelium trajectory, Red: nasal epithelium trajectory, Green: oral epithelium trajectory. **(E)** Dot plot shows enriched pathways (y-axis) of driver genes for each trajectory (x-axis). Dot is scaled by enrichment ratio and colored by significance. **(F)** Dot plot shows enriched motifs (y-axis) for each trajectory (x-axis). Dot is scaled by motif enrichment ratio and colored by significance. The position weight matrix of representative motifs Smad2::Smad3, Klf1, and Sox6 are included.

We next performed trajectory analysis of main subtypes as described above. As expected, the random walks mainly started from the undifferentiated epithelium and end in different terminal states (**Figure 6B, left panel**). We identified three trajectories, from undifferentiated epithelium to (1) dental epithelium, (2) nasal epithelium, and (3) oral epithelium (**Figure 6B, right panel**), validated by RNA velocity analysis (**Figure 6C**). We then identified driver genes for each trajectory, such as *Ccdc146* (correlation = 0.733, adjusted *p*-value < 2.2×10^-16^) for nasal epithelium trajectory, *Fras1* (correlation = 0.618, adjusted *p*-value < 2.2×10^-16^) for dental epithelium trajectory, and *Erbb4* (correlation = 0.697, adjusted *p*-value < 2.2×10^-16^) for oral epithelium trajectory (**Figure 6D**). We noticed that driver genes of the oral epithelium trajectory had negative fate correlations to the other two trajectories, such as *Nkain3* (correlationoral = 0.777, correlationnasal = −0.317, correlationdental = −0.605, adjusted *p*-values < 2.2×10^-16^) (**Figure 6D**). GSEA revealed enrichment of cilium related pathways (enrichment ratio = 15.13, adjusted *p*-value < 2.2×10^-16^) for nasal epithelium trajectory, odontogenesis (enrichment ratio = 6.14, adjusted *p*-value = 2.06×10^-4^) for dental epithelium trajectory, membrane development (enrichment ratio = 13.93, adjusted *p*-value = 7.24×10^-4^) for oral epithelium trajectory, respectively (**Figure 6E**). Subsequent motif enrichment analysis pinpointed the regulatory role of Smad2::Smad3 for dental epithelium trajectory (motif adjusted *p*-value = 3.32×10^-7^, *Smad3* gene *p*-value = 1.10×10^-7^, *Smad2* gene *p*-value = 0.13), Klf1 for nasal epithelium trajectory, and Sox6 for oral epithelium trajectory (**Figure 6F**).

## DISCUSSION

Dynamic gene expression patterns, driven by the dynamic activity of transcription factors (TFs) and accessibility of their binding sites [e.g., *cis*-regulatory elements (CREs)], underlie the formation of the secondary palate. Single-cell multi-omics technology generates paired gene expression and chromatin accessibility measurements, which paves the way for accurately tracking gene regulation dynamics. In this study, we generated time-series single-cell multi-omic datasets of the mouse secondary palate from E12.5 to E14.5 to dissect lineage-determining TFs that govern the developmental process. Our study is the first that profiled multiple modalities of developing mouse secondary palate within the same cells at single-cell resolution.

Cell type annotation is challenging due to the lack of reference datasets and the general rareness of data for secondary palate development. Automated cell type annotation approaches on scRNA-seq data, such as SingleR (Aran et al., 2019) and deCS (Pei et al., 2022), are not applicable. In this study, cell types were defined through extensive manual curation of marker genes, which were validated using an independent scRNA-seq dataset taken from a similar tissue (Han et al., 2021). We discovered subpopulations in CNC-derived mesenchymal cells that aligh with the *in vivo* anatomical locations, which were validated by our experiments, including *in situ* hybridization, quantitative RT-PCR, and bulk RNA sequencing. The established cell types and subtype-specific gene expression profiles, together with the chromatin accessibility profiles, can be used as the reference and facilitate cell type annotations in future analyses.

Pseudotemporal ordering of cells from time-series scRNA-seq datasets requires integration of time information, where traditional approaches are not applicable (Tran and Bader, 2020). In this study, we applied the Wadding-Optimal Transport algorithm (Schiebinger et al., 2019) to connect cells between adjacent time points and reconstructed five trajectories, demonstrating the continuous developmental landscape of cell states. We identified 556 and 586 driver genes for anterior and posterior palatal mesenchymal trajectories, respectively, with supporting evidence from previous publications. For example, *Meox2* was a driver gene for the posterior palatal trajectory and it was previously reported that *Meox2*-null and heterozygous knockout mice exhibited posterior cleft palate due to post-fusion breakdown of palatal shelves (Jin and Ding, 2006). *Msx1*, a driver gene for dental mesenchymal trajectory, was shown to regulate the cell proliferation of dental mesenchymal cells(Feng et al., 2013) and tooth morphogenesis (Chen et al., 1996). Runt-related transcription factor 2 (*Runx2*) is known to regulate tooth and bone formation during the differentiation of CNC-derived cells (James et al., 2006), which exhibited high fate correlations to both dental and osteoblast trajectories in our study.

By connecting open chromatin signals indicative of accessible CREs at the DNA level with gene expression at the RNA level, we identified a list of TFs that control each trajectory by binding to CREs to regulate the expression of the above-mentioned driver genes. Consistent with our results, *Shox2* null mice exhibited cleft palate that were confined to the anterior region (Hilliard et al., 2005; Yu et al., 2005). Importantly, the posterior palate in *Shox2* null mice was intact, underscoring the gene expression and regulation differences along the anterior-posterior axis of the palate. Another key regulator during the middle of the anterior palatal trajectory, Runx1, was reported to regulate the fusion and its deficiency caused cleft in the anterior palate (Charoenchaikorn et al., 2009; Sarper et al., 2018). Dlx1 and Dlx2 were identified as the top regulators for the posterior palatal mesenchymal trajectory. Concordant with our findings, a previous study showed that *Dlx1/2* double knockout mice developed cleft palate due to the vertical growth failure of posterior palatal shelves (Jeong et al., 2012).

Although our study revealed dynamic gene regulation programs, it has limitations in explaining the three-dimensional processes, such as the reorientation of palatal shelves. Therefore, it would be interesting to integrate with spatial information, such as 10x Genomics Visium technology that overlay gene expression data with the morphological context in tissues (Saltz et al., 2018), to reveal spatial expression patterns and elucidate the mechanisms of palatal elevation and reorientation at the molecular level. Single-cell proteome data can also be integrated to quantify the downstream protein levels during development (Brunner et al., 2022). Furthermore, *in vivo* validations in developing mouse embryos are needed to confirm the regulatory role of identified lineage-determining TFs, such as the knockout of specific TFs or lineage tracing experiments. Computation approaches that identify driving TFs in scRNA-seq data (Simon et al., 2020) can also be employed to validate the results. Future studies may involve the application of constructed landscapes to related diseases. For example, chromatin accessibility profiles of cell types can be used to train a deep learning model to predict noncoding mutations in cleft palate and prioritize *de novo* genetic variants.

Overall, we built a single-cell multi-omics atlas of the developing mouse secondary palate covering four developmental stages by simultaneously profiling gene expression and chromatin accessibility from the same cells. The optimal transport-based approach connected adjacent time points and recovered continuous landscapes during development. Constructed five trajectories represented continuous differentiation of CNC-derived multipotent mesenchymal cells to terminally differentiated subpopulations. By linking open chromatin signals to gene expression, we characterized a list of lineage-determining transcription factors, including Shox2 for the anterior palatal mesenchymal trajectory and Dlx1/2 for the posterior palatal mesenchymal trajectory. In conclusion, our study charted epigenetic and transcriptional dynamics during palatogenesis and provided a valuable resource for the community to facilitate future research of cleft palate.

## ACKNOWLEDGMENTS

This study was supported by grants from the National Institutes of Health (R01DE030122, R01DE029818 and R01LM012806). F.Y. is a CPRIT Predoctoral Fellow in the Biomedical Informatics, Genomics, and Translational Cancer Research Training Program (BIG-TCR) funded by Cancer Prevention & Research Institute of Texas (CPRIT RP210045). The sequencing data were generated by the UTHealth Cancer Genomics Core funded by CPRIT (RP180734). The funders had no role in the study design, data collection, analysis, decision to publish, or preparation of the manuscript.

## AUTHOR CONTRIBUTIONS

Conceptualization, Z.Z., J.I., and L.S.; Methodology, L.S., F.Y., C.I., J.I., and Z.Z.; Formal Analysis, F.Y., L.S., A.S., C.I., and G.P.; Resources: J.I., A.S., C.I., and Z.Z.; Data generation: J.I., C.I., H.Y., X.C., and M.Y.; Writing – Original Draft, F.Y., A.S., C.I., and X.C.; Writing – Review & Editing, Z.Z., L.M.S., J.I., A.S., C.I.; Visualization: F.Y., L.S., and C.I.; Funding Acquisition, Z.Z. and J.I.; Supervision, Z.Z., J.I., and L.S. All authors read and approved the final manuscript.

## STAR METHODS

### RESOURCE AVAILABILITY

#### Lead contact

Further information and requests for resources and analysis should be directed to and will be fulfilled by the lead contact, Dr. Zhongming Zhao.

#### Data and code availability

We are in process of data deposition at GEO and will update the accession number once finished. Previously published soft palate data that were reanalyzed in this study is available under accession codes GSE155928. All R and Python scripts supporting the findings of this paper are available upon request.

### EXPERIMENTAL MODEL AND SUBJECT DETAILS

#### Tissue preparation, dissociation, and nuclei extraction

Palatal shelves were isolated from time-mated C57BL/6J mice (000664, Jackson Laboratory) at E12.5, E13.5, E14.0, and E14.5. All mice were maintained in the animal facility of UTHealth under the 12 hours light/dark cycle and access to food/water ad libitum. The protocol was approved by the Animal Welfare Committee (AWC) and the Institutional Animal Care and Use Committee (IACUC) of UTHealth (AWC 19-0079).

Single-cell suspensions were prepared from pooled paired secondary palatal shelves of three embryos at E12.5, two embryos at E13.5, and one embryo at E14.0 and E14.5, respectively. The micro-dissected palatal shelves were treated with 0.25% trypsin and 0.05% EDTA (150 μL) for 5 min at 37 °C with gentle agitation (300 rpm). The dissociated cell mixtures were then suspended with 300 μL Dulbecco’s Modified Eagle’s Medium (DMEM, Millipore Sigma) supplemented with heat-inactivated 10% fetal bovine serum (FBS).The cells were centrifuged at 500 g for 5 min at 4 °C and the cell pellets were resuspended and incubated with chilled 300 μL of 0.1x lysis buffer [10 mM Tris-HCl pH 7.4, 10 mM NaCl, 3 mM MgCl_2_, 1% BSA, 1mM DTT, 1U/μL RNase inhibitor, 0.01% Tween-20, 0.01% Nonidet P40 Substitute, and 0.001 % Digitonin] for 3 min on ice, followed by stopped with chilled 300 μL wash buffer [10 mM Tris-HCl pH 7.4, 10 mM NaCl, 3 mM MgCl2, 1% BSA, 1 mM DTT, 1U/μL RNase inhibitor, and 0.1% Tween-20]. The cells were then collected by centrifuge at 500 g for 5 min at 4 °C, rinsed with 200 μL wash buffer twice, and re-suspended in Diluted Nuclei Buffer [1x Nuclei Resuspension Buffer, 1 mM DTT, and 1U/μL RNase inhibitor]. Isolated single-cell nuclei were filtered using a cell strainer (40 μm pore size) and inspected under a microscope to ensure they were successfully dissociated into single cell level for subsequent sequencing.

#### Single-cell multiome data generation

The single cell libraries were constructed by following the 10x Genomics Chromium Next GEM Single Cell Multiome ATAC + Gene Expression protocol (CG000338). Briefly, nuclei suspensions were incubated with a transposase, which fragmented the DNA in open regions of the chromatin and added the adapter sequences to the ends of the DNA fragments. The transposed nuclei were loaded onto Chromium Next GEM Chip J (PN-1000234, 10xGenomics, Pleasanton, CA) with partitioning oil and barcoded singlecell gel beads, followed by PCR amplification. The ATAC library and the gene expression library were then prepared separately. The quality of the libraries was examined using Agilent High Sensitive DNA Kit (#5067-4626) by Agilent Bioanalyzer 2100 (Agilent Technologies, Santa Clara, USA). The library concentrations were determined by qPCR using Collibri Library Quantification kit (#A38524500, Thermo Fisher Scientific) on a QuantStudio3 (ThermoFisher Scientific). We then pooled the libraries evenly and performed the paired-end sequencing on an Illumina NextSeq 550 System (Illumina, Inc., USA) using High Output Kit v2.5 (#20024907, Illumina, Inc., USA).

### METHOD DETAILS

#### Single-cell multiome data processing

The 10x Genomics Cell Ranger ARC (v2.0.0) pipeline was used to process the multiome data. Raw sequencing data were first converted to fastq format using ‘cellranger-arc mkfastq’. The raw files of RNA-seq and ATAC-seq libraries from the same sample were aligned to the UCSC mouse genome (mm10) and quantified using ‘cellranger-arc count’. Samples were aggregated using ‘cellranger-arc aggr’ to normalize the sequencing depth.

The raw RNA count matrix and ATAC fragment data were further processed using R packages Seurat (v4.0.3) (Hao et al., 2021) and Signac(v1.5.0) (Stuart et al., 2021), respectively. Filtering based on RNA-assay metrics (200< nCount_RNA < 100,000, nFeature_RNA < 7,500, percent.mt < 20) and ATAC-assay metrics (200 < nCount_ATAC < 100,000, nucleosome_signal < 2, TSS.enrichment > 1) resulted in 37,329 cells. The average depth is 73,521 reads per cell, yielding an average of 2,472 genes per cell.

The gene expression count matrix was normalized using SCTransform. Principal component (PC) analysis was based on the top 3,000 highly variable features. Uniform Manifold Approximation and Projection (UMAP) visualization was constructed using the first 30 PCs.

For the ATAC data, peak calling was performed using MACS2 package (Zhang et al., 2008) with *CallPeaks* function in Signac (version1.5.0). Peaks that overlapped with genomic blacklist regions for the mm10 genome were removed (Amemiya et al., 2019). Each peak represents one potential *cis*-regulatory DNA element (CRE). The CRE count matrix was then normalized using Latent Semantic Indexing (LSI), including term-frequency (TF) inverse-document frequency (IDF), and Singular value decomposition (SVD). The first LSI component is removed from the downstream analysis as it was highly correlated with sequencing depth.

The gene activity was quantified using *GeneActivity* function in Signac (version1.5.0), which aggregated chromatin accessibility intersecting the gene body and promoter regions.

#### Projection of external scRNA-seq dataset onto our scRNA-seq manifold

To validate the annotated major cell types, we downloaded a scRNA-seq dataset of mouse soft palate, which is the posterior third of the palate, from a recent publication (Han et al., 2021). SCTransform normalization was conducted, followed by PC analysis. The first 30 PCs were used to find anchors between these two datasets using *FindTransferAnchors* function in the Seurat package. The *RunUMAP* function was used to calculate the UMAP coordinates of our dataset with parameters stored in the object (return.model = TRUE). The *MapQuery* function was then used to calculate the coordinates of the external dataset using the same ‘uwot’ model parameters.

#### In-situ hybridization

The E14.5 C57BL/6J mouse embryos (n=6) were dissected from a time-pregnant mother and fixed in 4% paraformaldehyde overnight at 4 °C, dehydrated in a graded ethanol series, and embedded in paraffin. Paraffin sections were cut at 4 μm thickness under RNase-free conditions. *In situ* hybridization was performed using the RNAscope 2.5 Assay platform (ACD, 322360) using specific probes for *Cyp26b1* (ACD, 454241), *Efnb2* (ACD, 477671), *Inhba* (ACD, 455871), *Meox2* (ACD, 823191), *Nrp1* (ACD, 471621), *Prickle1* (ACD, 832641), *Satb2* (ACD, 413261), *Shox2* (ACD, 579051), *Sim2* (ACD, 1110401), and *Trps1* (ACD, 879001). The color images were obtained under a light microscope (BX43, Olympus).

#### Quantitative RT-PCR

The anterior (n=6) and posterior (n=6) palatal shelves were microdissected from E14.5 C57BL/6J mouse embryos for qRT-PCR. Total RNAs isolated from each region were collected using the QIAshredder and RNeasy mini extraction kit (QIAGEN), as previously described (Suzuki et al., 2015). *Gapdh* was used as an internal housekeeping gene control. The PCR primers used in this study are listed in **Table S2**.

#### Bulk RNA-seq analysis

The anterior (n=3) and posterior (n=3) palatal shelves of E14.0 C57BL/6J mice were isolated and subjected to bulk RNA sequencing. The raw sequenced files were mapped to the mouse reference genome mm10 using HISAT2 (Zhang et al., 2021). StringTie (Shumate et al., 2022) was used to quantify the counts. We then used R package DESeq2 (version 1.30.1) to perform the differential gene expression tests (Love et al., 2014). To project the bulk RNA-seq data into the scRNA principal component space, count matrices from both datasets were first integrated and normalized using *voom* function in the R package limma (version 3.46.0). We performed PCA independently using normalized scRNA data. The normalized bulk RNA-seq data were then projected into the scRNA space using identical principal component gene loadings.

#### CRE-gene linkage analysis

We identified CRE-to-gene links using *LinkPeaks* function in Signac (Stuart et al., 2021) based on the approach originally described by SHARE-seq (Ma et al., 2020). The Pearson correlation coefficient was calculated between gene expression and CRE accessibility. Only CREs within a certain distance (bp) from the gene TSS were included in the model (default: 5×10^5^). The GC content, overall accessibility, and CRE size were included in the model as covariates to correct the bias. Benjamini-Hochberg method was used to adjust *p*-values (Benjamini and Hochberg, 1995). Only high-confidence CRE-gene links with adjusted *p*-value < 0.05 and coefficients > 0 were retained for downstream analysis.

#### DNA sequence motif enrichment analysis

A total of 196 position weight matrices for *Mus musculus* (species code 10090) were loaded from the JASPAR 2020 database (Fornes et al., 2020) using *getMatrixSet* function in TFBSTools package (version 1.32.0). For a set of differentially accessible CREs, we applied *FindMotifs* function with default parameters to find enriched motifs. Meanwhile, to facilitate the visualization of motif activity, we calculated the motif activity matrix using ChromVAR (version 1.16.0) (Schep et al., 2017).

#### WOT-based terminal state likelihood analysis

The Wadding-Optimal Transport (WOT) was employed to reconstruct the trajectories (Schiebinger et al., 2019). Specifically, for a cell at time *ti*, WOT traced its most likely ancestors and descendants to recover the trajectories by calculating the transition probabilities to cells at time *ti+1* and *ti-1*. We imported *WOTKernel* from *cellrank.external.kernels* for the following analysis (Lange et al., 2022). The growth rates were estimated using the predefined gene proliferation set. The cell-cell transition matrix between adjacent time-points was then calculated using *compute_transition_matrix* function with default parameters (growth_iters=3, growth_rate_key=“growth_rate_init”, last_time_point=“connectivities”). The random walks were simulated (n_sims=300). To compute the macrostates, a Generalized Perron Cluster Cluster Analysis (GPCCA) estimator was initialized with WOT connectivity kernel(Reuter et al., 2018). The inferred macrostates were set as terminal states of five trajectories. The fate probabilities to each terminal state were computed per cell using *compute_absorption_probabilities* function with default parameters (solver=“gmres”). To identify driver genes, we computed the correlation between the fate probabilities and gene expression for each trajectory using *compute_lineage_drivers* function. Multiple testing correction was controlled by the Benjamini-Hochberg method (Benjamini and Hochberg, 1995). Only the genes with adjusted *p*-values less than 0.05 and correlations greater than 0.05 were considered driver genes.

#### Diffusion pseudotime estimation

We used the built-in function of the Python package Scanpy (version 1.9.1) to estimate the diffusion pseudotime(Haghverdi et al., 2016; Wolf et al., 2018). Specifically, the raw count matrix was loaded in as an AnnData object, followed by standard preprocessing. The neighborhood graph was calculated using *sc.pp.neighbors* function with default parameters (random_state=0). We specified a random cell in CNC-derived multipotent cells (cluster 5) as the root cell. The diffusion pseudotime was then estimated using *scanpy.tl.dpt* function.

#### RNA velocity and CellRank analysis

The possorted bam files from Cellranger output were passed to velocyto (version: 0.17.15) (La Manno et al., 2018) to estimate the RNA velocities of single cells. The generated loom file contained data matrices of spliced and unspliced reads and was further processed by scVelo (version 0.2.4)(Bergen et al., 2020). Seurat-processed gene expression count matrix and UMAP coordinates were converted to “AnnData” object and merged with the velocyto-derived object using *scVelo.utils.merge* function. The merged dataset was filtered using the *scVelo.pp.filter_and_normalize* with default parameters (min_shared_counts = 10, n_top_genes = 2,000) and the moments were computed using *scVelo.pp.moments*. The velocity was then calculated using *scVelo.tl.velocity* (mode=stochastic). The estimated velocity vectors were projected and visualized in previously calculated embedding.

The initial and terminal state likelihood based on RNA velocity information was estimated using *cellrank.tl.terminal_states* and *cellrank.tl.initial_states* functions with default parameters (weight_connectivities=0.3).

#### Trajectory analysis

To identify how and when driver genes were expressed and regulated along each trajectory, we extracted cells with high fate probabilities (fate probability > 75% quantile). The extracted cells were then ordered by diffusion pseudotime. The driver genes were cut into three groups based on quantiles. For each group of driver genes, we conducted gene set enrichment analysis using the R package WebGestalt (version 0.4.4) (Liao et al., 2019). The non-redundant Gene Ontology (GO) Biological Process terms were used for pathway annotations. The minimum number of genes in the pathways was set to 5 and the maximum was set to 500. The Benjamini-Hochberg method was used to adjust *p*-values (Benjamini and Hochberg, 1995). Those pathways with adjusted *p*-values less than 0.05 were considered statistically enriched. We also performed CRE-gene linkage analysis and motif enrichment analysis as described above.

## Supplemental information

Supplemental figures and tables can be found in separated pdf file.

---

**Supplemental Figure 1.**
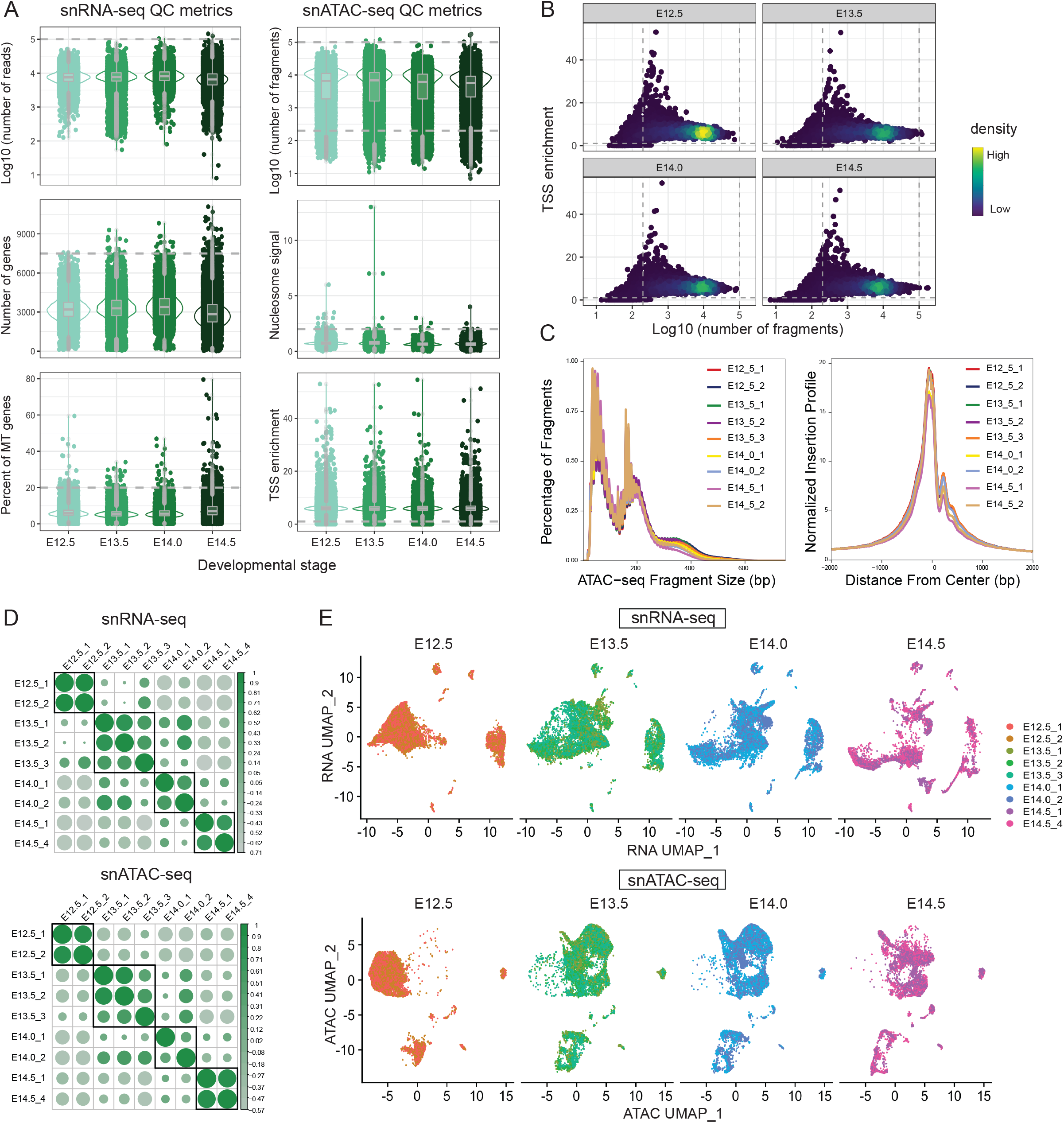
Data quality of scATAC-seq and scRNA-seq libraries. **(A)** Violin plot show the distribution of (left) scRNA-seq and (right) scATAC-seq quality metrics, including the number of reads, number of genes, and mitochondrial (MT) gene fraction per cell, the number of fragments, nucleosome signal, and transcription start site (TSS) enrichment per cell in each developmental stage. E: embryonic day. **(B)** Scatter plot show scATAC-seq cell thresholding on TSS enrichment (y-axis) and the number of fragments (x-axis). Dashed lines represent filtering thresholds. Color represent density. **(C)** Top: Plot shows the normalized fragment count in each positive rekative to TSS (bp) for each sample. Bottom: Density plot show the distribution of fragment size for each sample. **(D)** Correlogram shows the correlation between samples in RNA assay (top) or ATAC assay (bottom). **(E)** UMAP visualization based on RNA assay (top) or ATAC assay (bottom) represent cells colored by each sample and split by developmental stage.

**Supplemental Figure 2.**
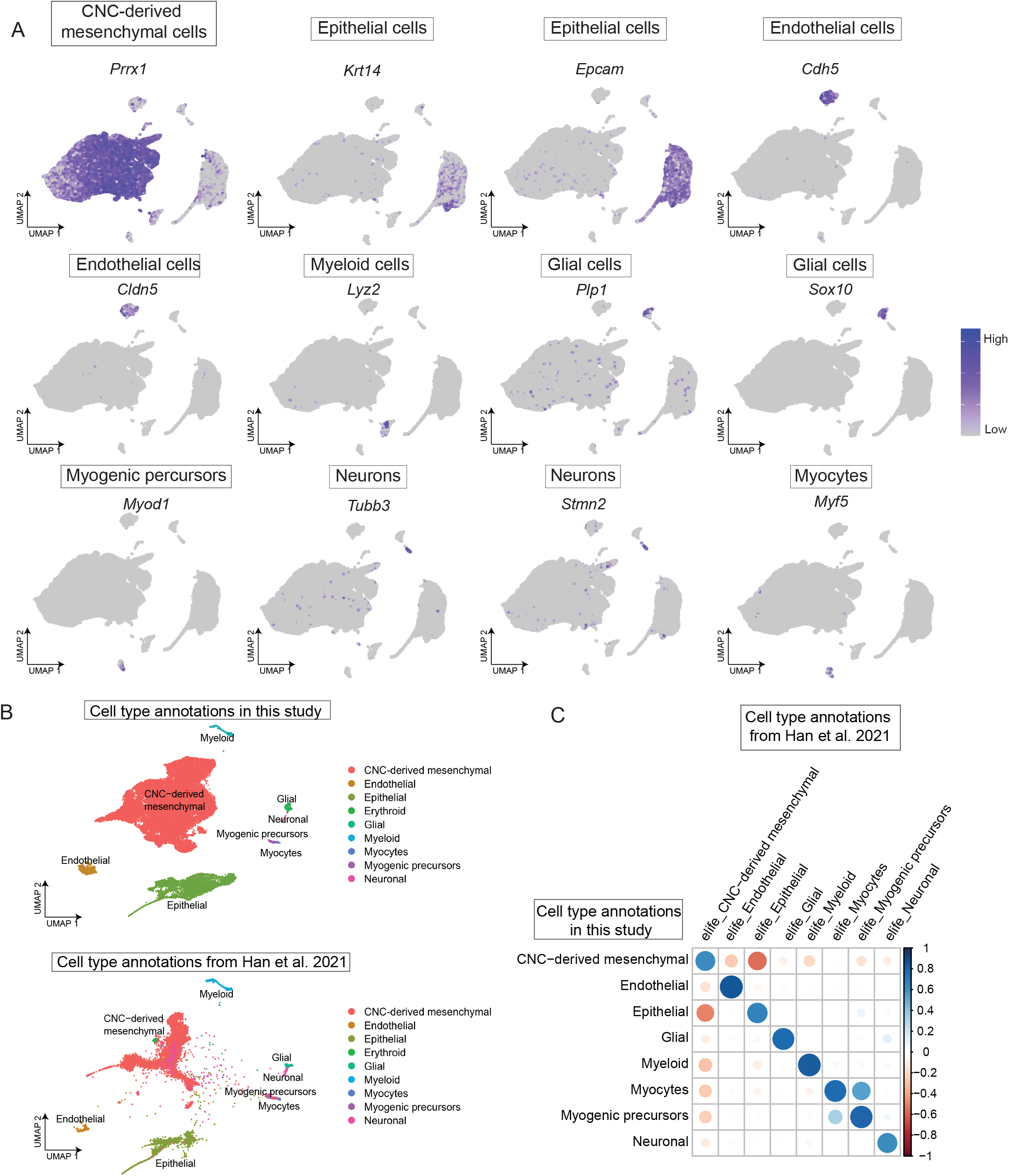
Comparison with external scRNA-seq dataset validate our cell type annotations. **(A)** Feature plot show marker gene expression in each major cell type. Marker genes and cell types were annotated on top of each plot. **(B)** Projection of external data from Han et al. 2021 into our snRNA-seq manifold, showing alignment of major cell types. **(C)** Correlogram shows the correlation between cell type annotations in our dataset (row) and annotations from Han et al. 2021 (columns). Each dot represents the correlation between one paired cell type and is colored from low (red) to high (blue).

**Supplemental Figure 3.**
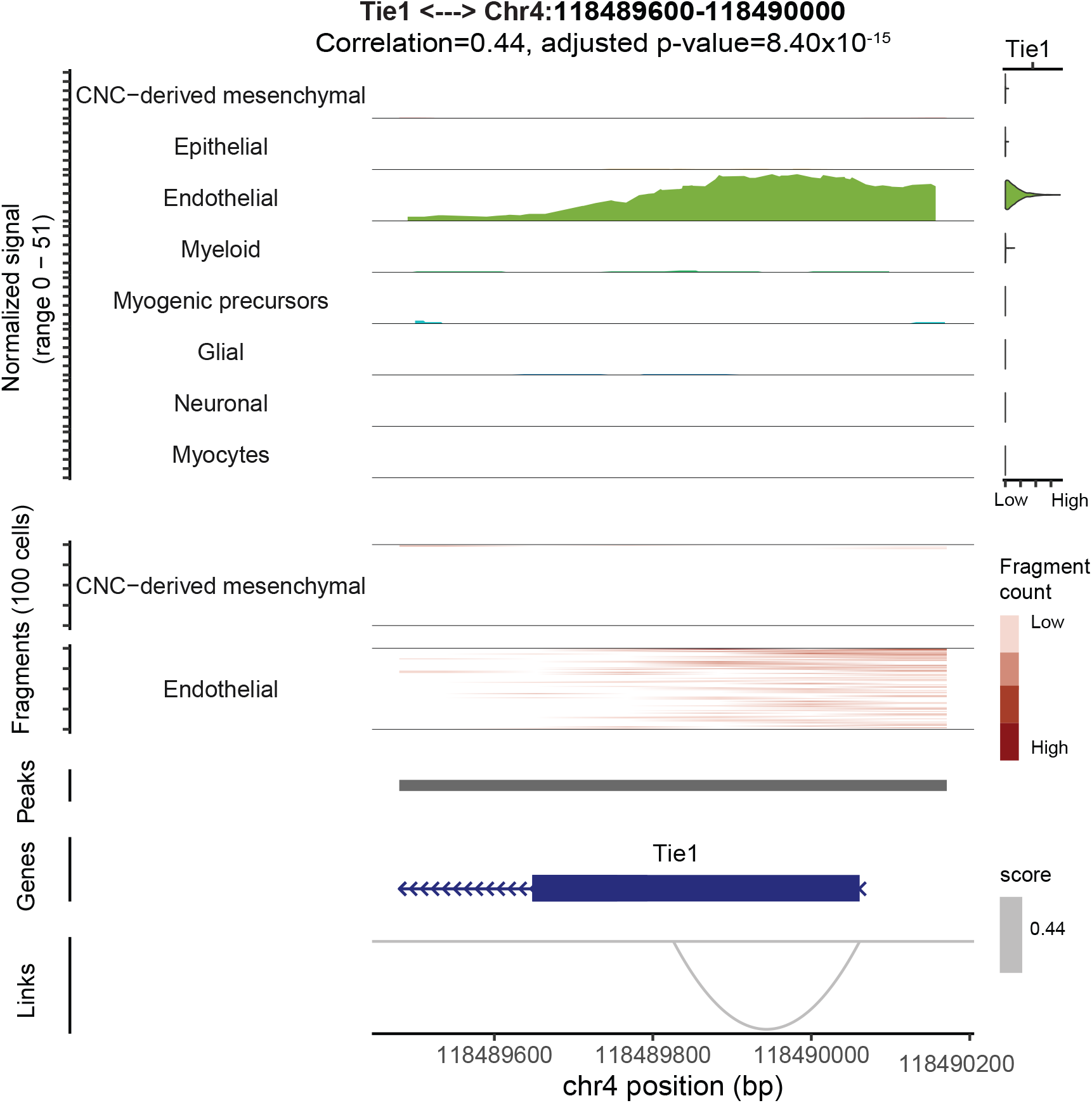
Visualization displaying positively peak-gene linkage specifically in Endothelial cells. Left: Genome Browser visualization of aggregated chromatic accessibility at the chr4:118489600-118490000 locus for each major cell type, coupled with Tie1 gene expression. Arcs at the bottom denotes linkage between Tie1 and Chr4:118489480-118490171.

**Supplemental Figure 4.**
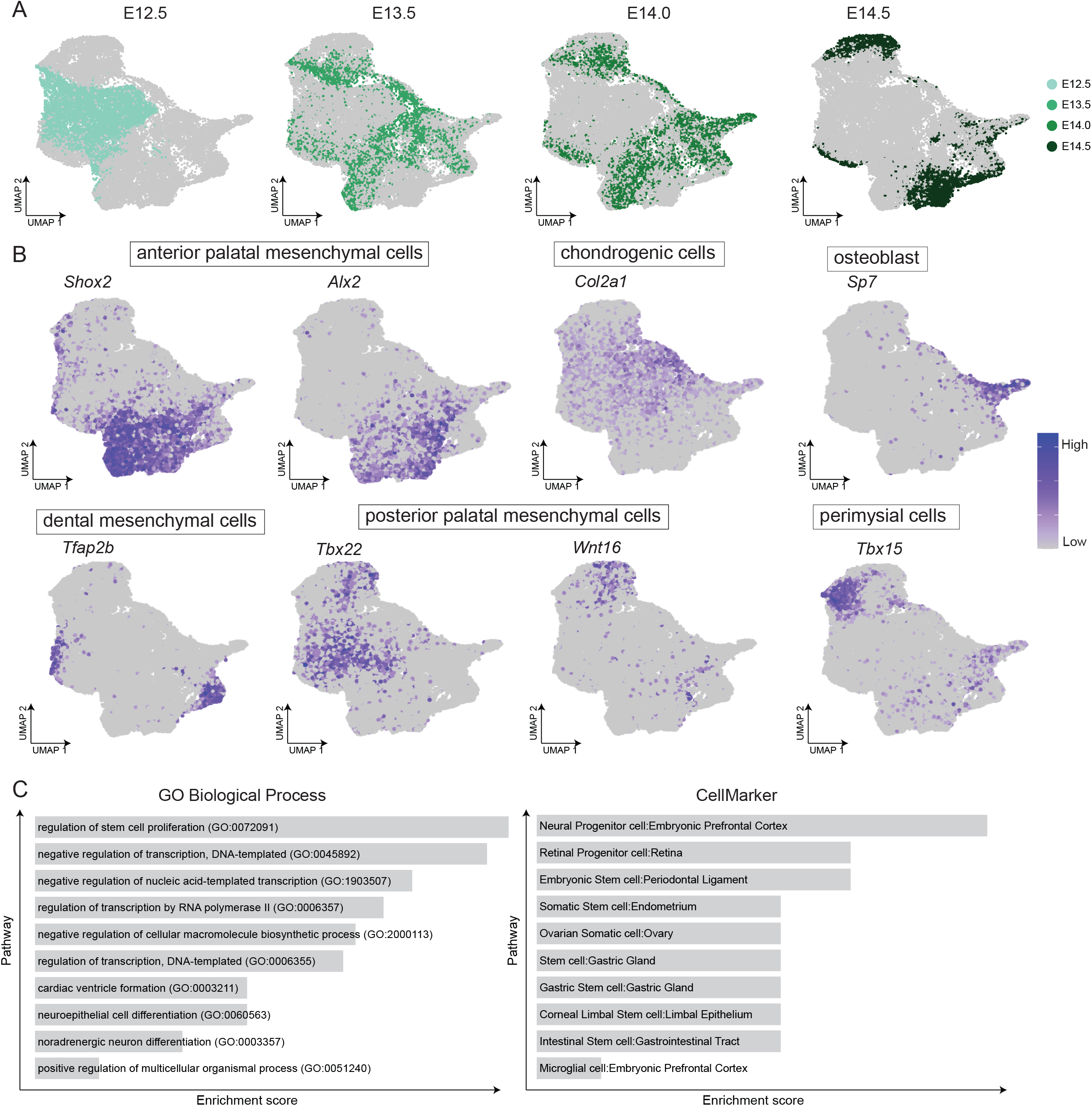
Cell subtype annotations for CNC-derived mesenchymal cells. **(A)** UMAP visualization corlored by developmental stage. (B) Feature plot of marker genes of each subtype of CNC-derived mesenchymal cells. **(C)** Barplot shows enriched gene ontology (GO) Biological Process pathways (left) and cell types (right) for cluster 5.

**Supplemental Figure 5.**
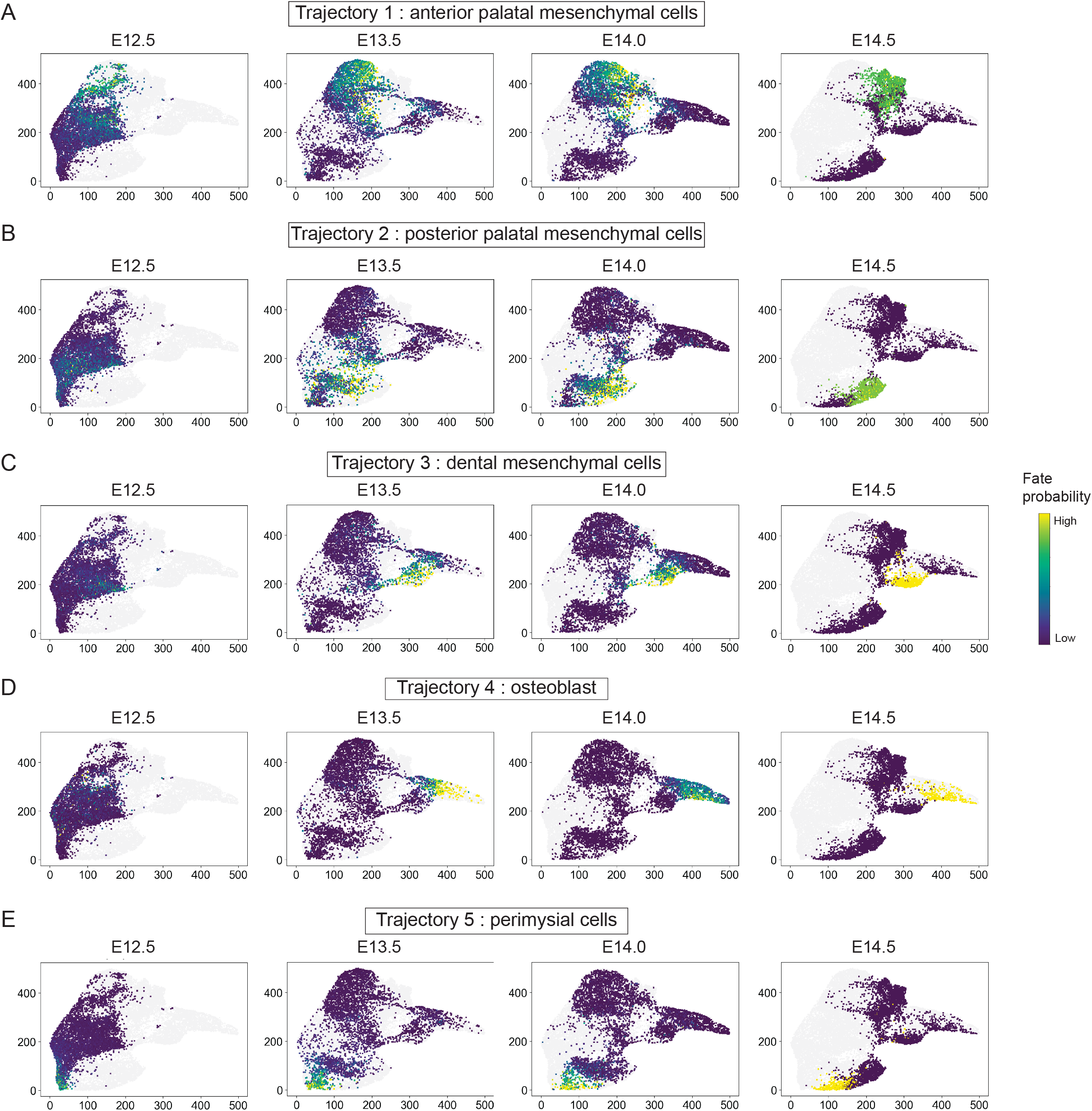
Trajectory analysis revealed the continuous differentiation of CNC-derived multipotent cells into different terminal states. **(A)** Trajectory 1: anterior palatal mesenchymal cells, **(B)** Trajectory 2: posterior palatal mesenchymal cells, **(C)** Trajectory 3: dental mesenchymal cells, **(D)** Trajectory 4: osteoblast, **(E)** Trajectory 5: perimysial cells. Each dot represent a cell and is colored by fate probabilities to each trajctory (high: yellow, low: dark brown).

**Supplemental Figure 6.**
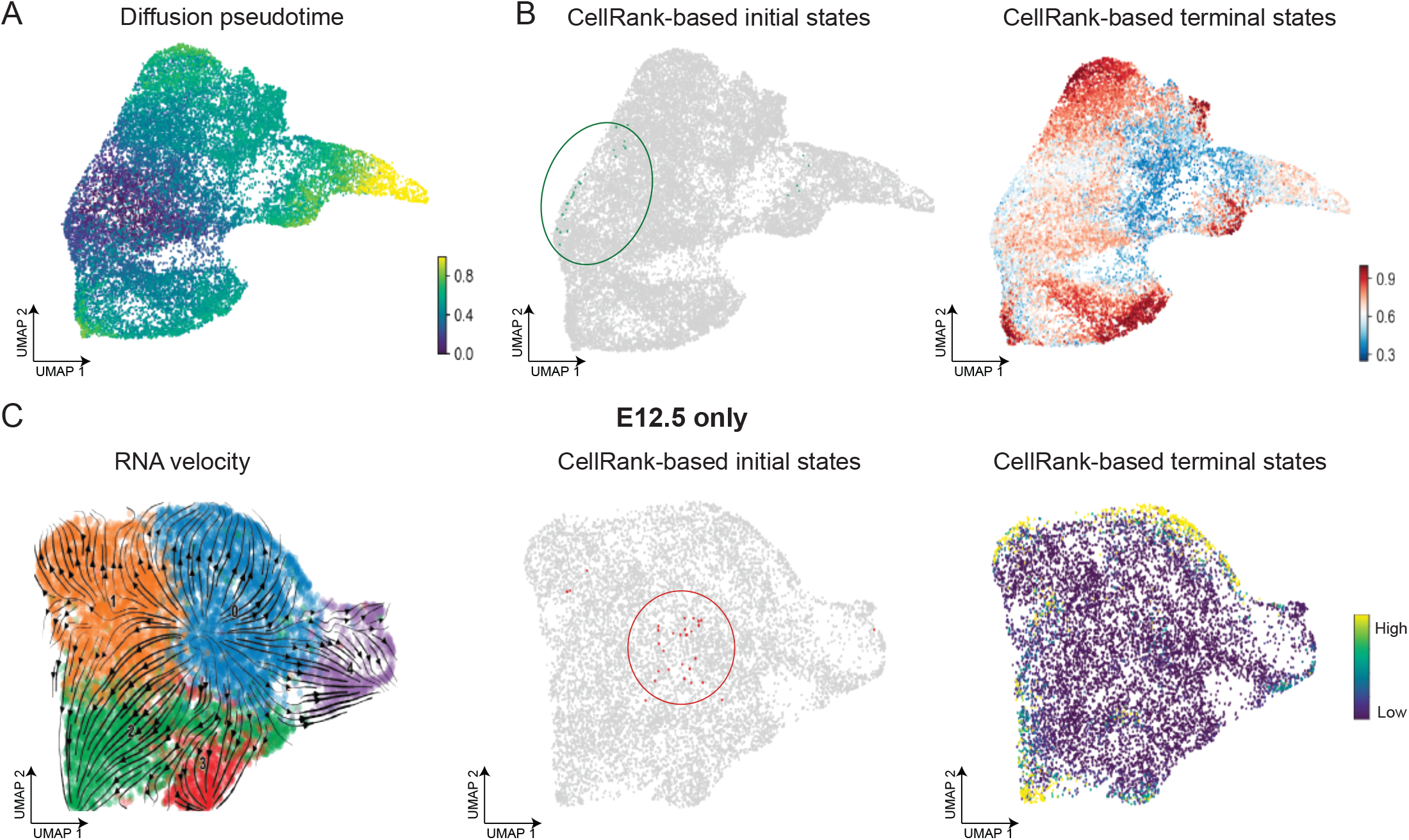
CellRank based on RNA velocity validated the inferred trajectories. **(A)** UMAP visualization shows inferred diffusion pseudotime. **(B)** UMAP visualization shows (left) CellRank-derived initial stages and (right) terminals states. **(C)** UMAP visualization of CNC-derived mesenchymal cells from E12.5 only, colored with (left) RNA velocity information, (middle) CellRank derived initial states, and (right) CellRank derived terminal states.

**Supplemental Figure 7.**
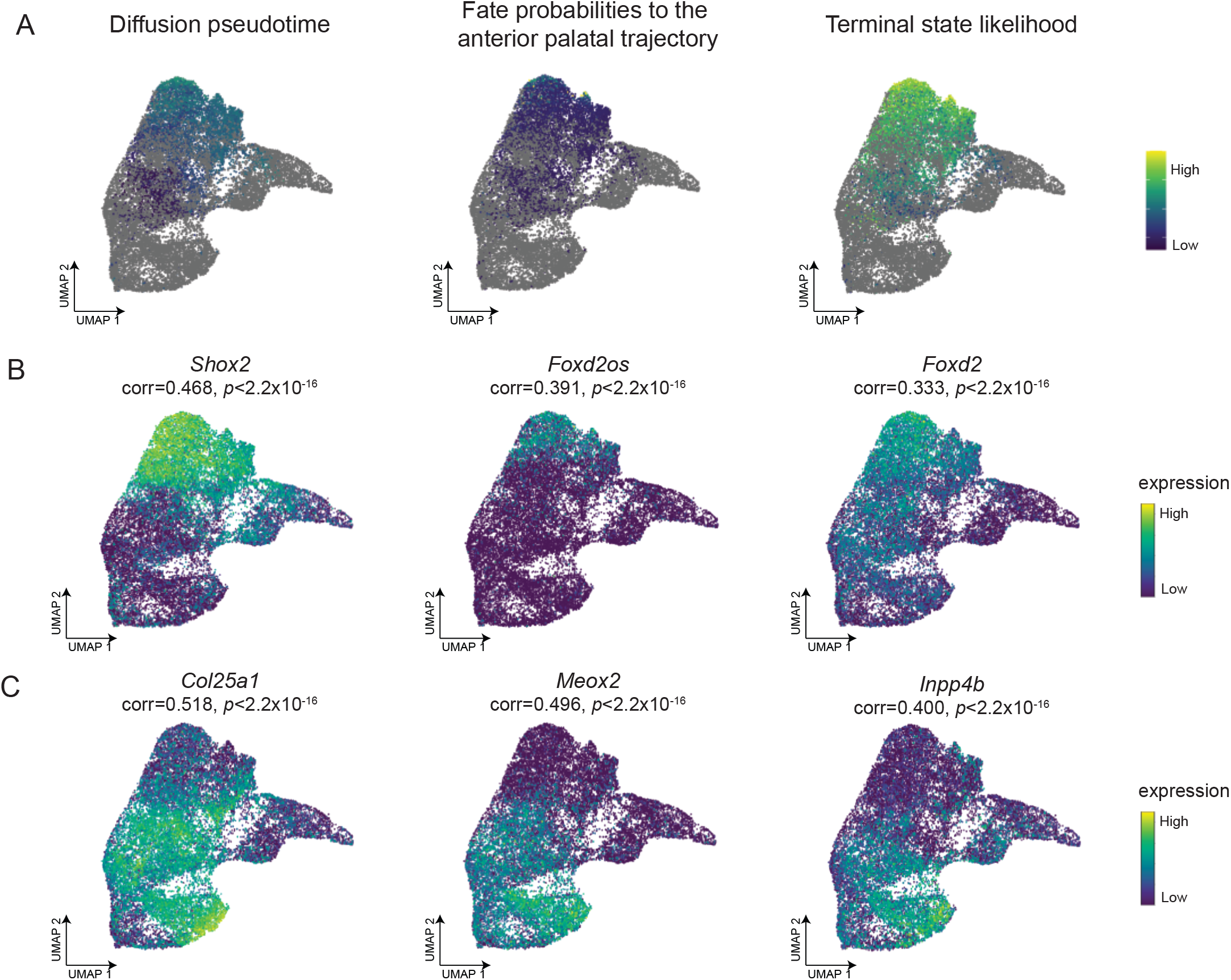
Driver genes were identified for each trajectory. **(A)** UMAP visualization showing anterior trajectory, colored by (left) diffusion pseudotime, (middle) fate probabilities to anterior trajectory, and (right) terminal state likelihood. **(B-C)** Feature plot show expression pattern of top driver genes for **(B)** anterior and **(C)** posterior trajectory. Each dot represent a cell and is colored by gene expression (yellow: high, dark brown: low). Gene name, adjusted p-value, and correlation with fate probabilities was annotated on top of each plot.

**Supplemental Table 1.** Quality control metrics for ATAC and GEX libraries in each sample.

| Library | ATAC Per Library Metrics |  |  | GEX Per Library Metrics |  |  | # of cells<br>before<br>preprocessi<br>ng | # of cells<br>after<br>preprocess<br>ing |
| --- | --- | --- | --- | --- | --- | --- | --- | --- |
|  | Total # of<br>read pairs | # of read<br>pairs per<br>cell | % of<br>reads<br>kept | Total # of<br>read pairs | # of read<br>pairs per<br>cell | % of<br>reads<br>kept |  |  |
| E12_5_1 | 253,080,554 | 37,571 | 52.90% | 175,630,078 | 26,073 | 90.00% | 6,736 | 6,348 |
| E12_5_2 | 201,627,408 | 34,267 | 61.40% | 144,278,831 | 24,520 | 100.00% | 5,884 | 5,498 |
| E13_5_1 | 189,257,170 | 86,616 | 80.60% | 176,962,262 | 80,989 | 41.50% | 2,185 | 1,693 |
| E13_5_2 | 217,328,102 | 89,141 | 41.00% | 156,982,892 | 64,390 | 46.60% | 2,438 | 2,226 |
| E13_5_3 | 209,954,979 | 43,397 | 55.40% | 174,273,262 | 36,021 | 70.50% | 4,838 | 4,527 |
| E14_0_1 | 194,201,004 | 40,602 | 100.00<br>% | 156,955,815 | 32,815 | 96.30% | 4,783 | 4,496 |
| E14_0_2 | 232,945,189 | 53,342 | 72.50% | 168,557,694 | 38,598 | 69.40% | 4,367 | 4,112 |
| E14_5_1 | 173,222,784 | 37,388 | 90.70% | 192,325,131 | 41,512 | 57.80% | 4,633 | 3,889 |
| E14_5_4 | 206,644,325 | 54,308 | 45.50% | 153,505,067 | 40,342 | 65.70% | 3,805 | 3,365 |

#: number, %: percent

**Supplemental Table 2.**
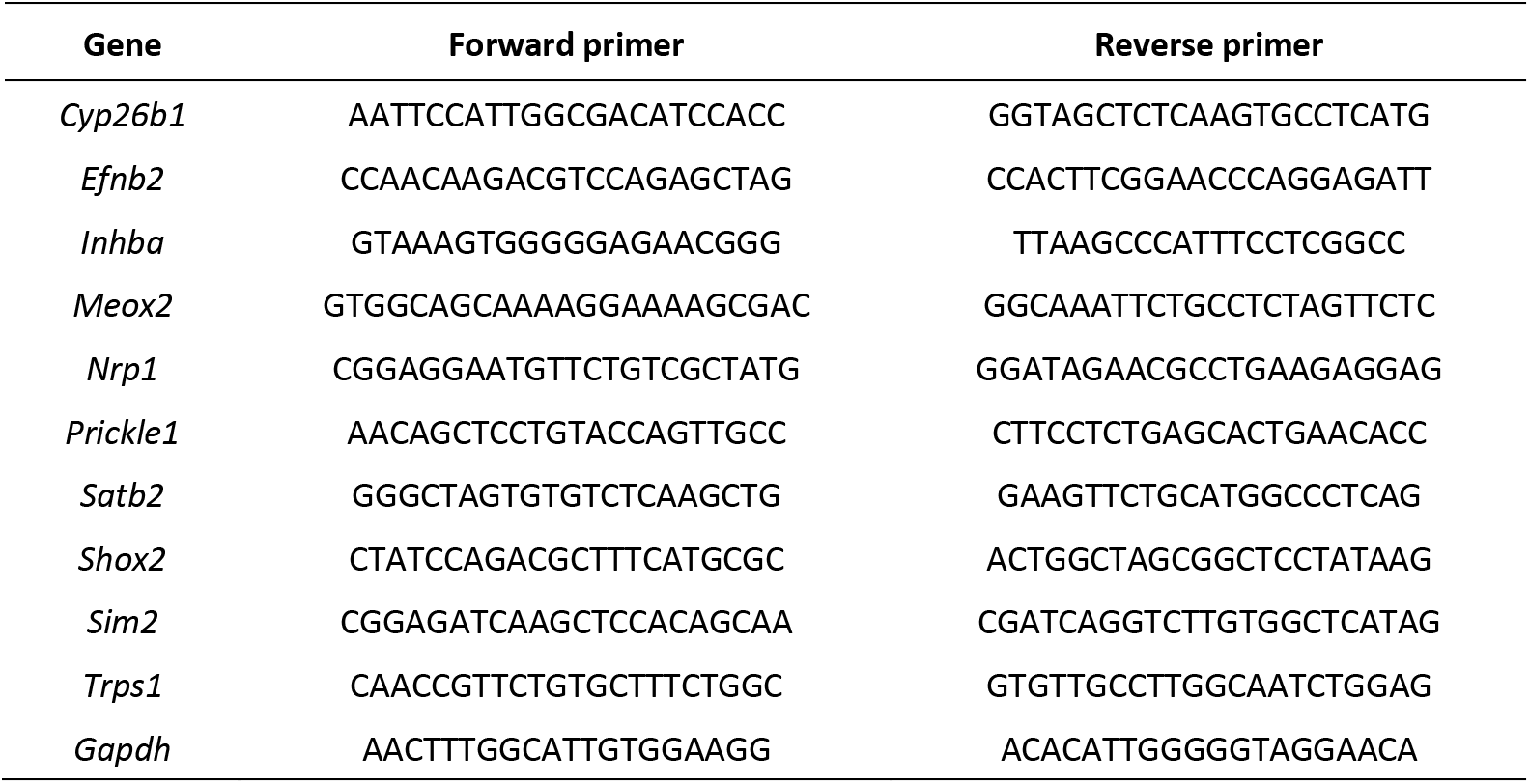
PCR primers for quantitative RT-PCR experiments.

